# The combination of stimulus-driven and modulatory inputs in visual thalamus depend on visual responsiveness and stimulus type

**DOI:** 10.1101/2023.10.18.562960

**Authors:** Lisa Schmors, Ann H. Kotkat, Yannik Bauer, Ziwei Huang, Davide Crombie, Lukas S. Meyerolbersleben, Sacha Sokoloski, Philipp Berens, Laura Busse

## Abstract

In the dorsolateral geniculate nucleus (dLGN) of the thalamus, stimulus-driven signals are combined with modulatory inputs such as corticothalamic (CT) feedback and behavioural state. How these shape dLGN activity remains an open question. We recorded extracellular responses in dLGN of awake mice to a movie stimulus, while photosuppressing CT feedback, and tracking locomotion and pupil size. To assess the relative impact of stimulus and modulatory inputs, we fit single neuron responses with generalized linear models. While including CT feedback and behavioural state as predictors significantly improved the model’s overall performance, the improvement was especially pronounced for a subpopulation of neurons poorly responsive to the movie stimulus. In addition, the observed impact of CT feedback was faster and more prevalent in the absence of a patterned visual stimulus. Finally, for neurons that were sensitive to CT feedback, visual stimuli could be more easily discriminated based on spiking activity when CT feedback was suppressed. Together, these results show that effects of modulatory inputs in dLGN depend on visual responsiveness and stimulus type.

## Introduction

Visual information is processed through a hierarchy of brain areas, which are connected by feedforward and feedback projections. Early in this hierarchy is the dorsolateral geniculate nucleus (dLGN) of the thalamus, a central node for visual information *en route* from the retina to the primary visual cortex (V1) (*1, 2*). The dLGN has long been recognized as one of the first visual stages that combines stimulus-driven inputs with additional modulatory inputs (*3*), such as signals arising from L6 cortico-thalamic (CT) feedback (*4, 5, 6, 7, 8*), and signals from the brainstem carrying information related to behavioural state (*9, 10, 11, 12*) and arousal (*10, 13, 14, 3*). Since these factors have often been investigated in separate studies, their combined effects on dLGN responses remain poorly understood. Overall, we lack a quantitative understanding of the relative strengths of these modulatory effects on dLGN responses during wakefulness and how they might interact and depend on stimulus-driven input.

On the one hand, it has been firmly established that, even during wakefulness, dLGN responses are modulated according to the animal’s internal (*15, 9*) and overt behavioural state (*16, 15, 17*). For instance, in the mouse, locomotion-(*11, 18*) or pupil-indexed arousal (*10, 9, 14, 19, 13*) are associated with overall enhancements of firing rates in dLGN, which seem to preferentially affect specific neuronal populations depending on their spatio-temporal feature selectivity (*18, 19*). Similar to related findings in the somatosensory system (*20*), the increase in dLGN firing rates seems to be a necessary condition for the sustained depolarisation of primary visual cortex during active behaviours (*9*).

On the other hand, little consensus has been achieved for dLGN modulations by CT feedback, where a plethora of previous studies have together highlighted its diverse and potentially stimulus-dependent effects. For instance, given that CT feedback can sharpen spatial dLGN RFs and increase contextual effects (*21, 22, 23, 6, 24, 7*), CT feedback might enhance or suppress overall firing rates, depending on the size, feature-selectivity and retinotopic position of dLGN neurons (*6, 7, 22*). The combination of enhancing and suppressing effects of CT feedback are likely mediated by a differential engagement of both direct excitatory and indirect inhibitory pathways, whose balance will depend on the stimulus selectivity, connectivity and intrinsic properties of corticothalamic neurons in L6 V1, neurons in the thalamic reticular nucleus (TRN) and dLGN (*8, 25*).

Generalized linear models (GLMs) provide an established framework for statistical analysis of neural responses, that can help to disentangle the combined impact of multiple stimulus-driven and modulatory influences and investigate their properties (*26, 27, 28*). While GLMs are relatively simple phenomenological models, they offer the advantage of being interpretable: for instance, GLM kernels learned for the visual stimulus approximate the integration by the spatio-temporal receptive field (RF), and kernels learned for any additional inputs represent spike-induced gain adjustments (*29, 27*). First applied in the retina (*30, 27*), GLMs have since then been used in numerous studies to separate influences of the visual stimulus and other variables, like spike history, interneuronal interaction effects, task-engagement, learning, reward prediction, task-related motor action, locomotion, and arousal (*31, 32, 33, 34, 35, 36, 37*). One recent extension of classical GLMs is to estimate RFs by choosing a set of cubic spline basis functions in order to encode smoothness, decrease the number of parameters, and thus be more data efficient (*38*).

Here, we investigated how feedforward, stimulus-driven signals, feedback signals, and behavioural state jointly influence dLGN activity in awake, head-fixed mice viewing a rich movie stimulus. We simultaneously recorded extracellular dLGN activity, mouse run speed and pupil size, while photosuppressing CT feedback. We then fitted a spline-GLM model containing kernels for the spatio-temporal RF, CT feedback and behaviour to predict responses of dLGN neurons. The learned kernels were biologically plausible, including diverse spatio-temporal RFs, as well as kernels for behaviour and CT feedback. We found that our model could successfully capture response components derived from the movie input, yielding RF kernels which were generally aligned with response features obtained from simple luminance steps. Including modulatory inputs overall improved the prediction of dLGN responses; the improvements, however, were most prominent for a subpopulation of neurons that were poorly predicted by the movie stimulus. Focusing on effects of CT feedback, we found that these effects depended on stimulus type in both the model and the data, being stronger, more prevalent and faster during the absence of a patterned visual stimulus. Finally, we used the spline-GLM for *in silico* experiments isolating the impact of CT feedback suppression, and demonstrate that stimulus discrimination for CT feedback-modulated neurons was enhanced. We conclude that effects of modulatory inputs in dLGN depend on visual responsiveness and stimulus type.

## Results

### dLGN responses to movies are modulated by behavioural variables and CT feedback suppression

To investigate how CT feedback, locomotion and arousal modulate thalamic responses, we recorded *in vivo* extracellular dLGN activity in response to a rich movie stimulus in four head-fixed mice together with running speed and pupil size, while randomly photo-suppressing CT feedback (**Figure 1a**). For photo-suppression of CT feedback, we conditionally expressed the soma-targeting, chloride-conducting channelrhodopsin stGtACR2-RFP (*39*) in L6 CT pyramidal cells, by injecting a small volume of Cre-dependent AAV into V1 of Ntsr1-Cre mice (*40*). The localisation of stGtACR2 to L6 CT somata and the accurate placement of electrodes were confirmed through post-mortem histological analyses (**Figure 1b**). During electrophysiological recordings, the mouse viewed a rich movie stimulus that consisted of a sequence of black-and-white clips from various feature films (‘movies’, **Figure 1c, top**). Here, photo-suppression occurred with 50% probability in each 1 second time bin. We also measured the mouse’s run speed and pupil size to infer the animal’s changing behavioural state.

**Figure 1.**
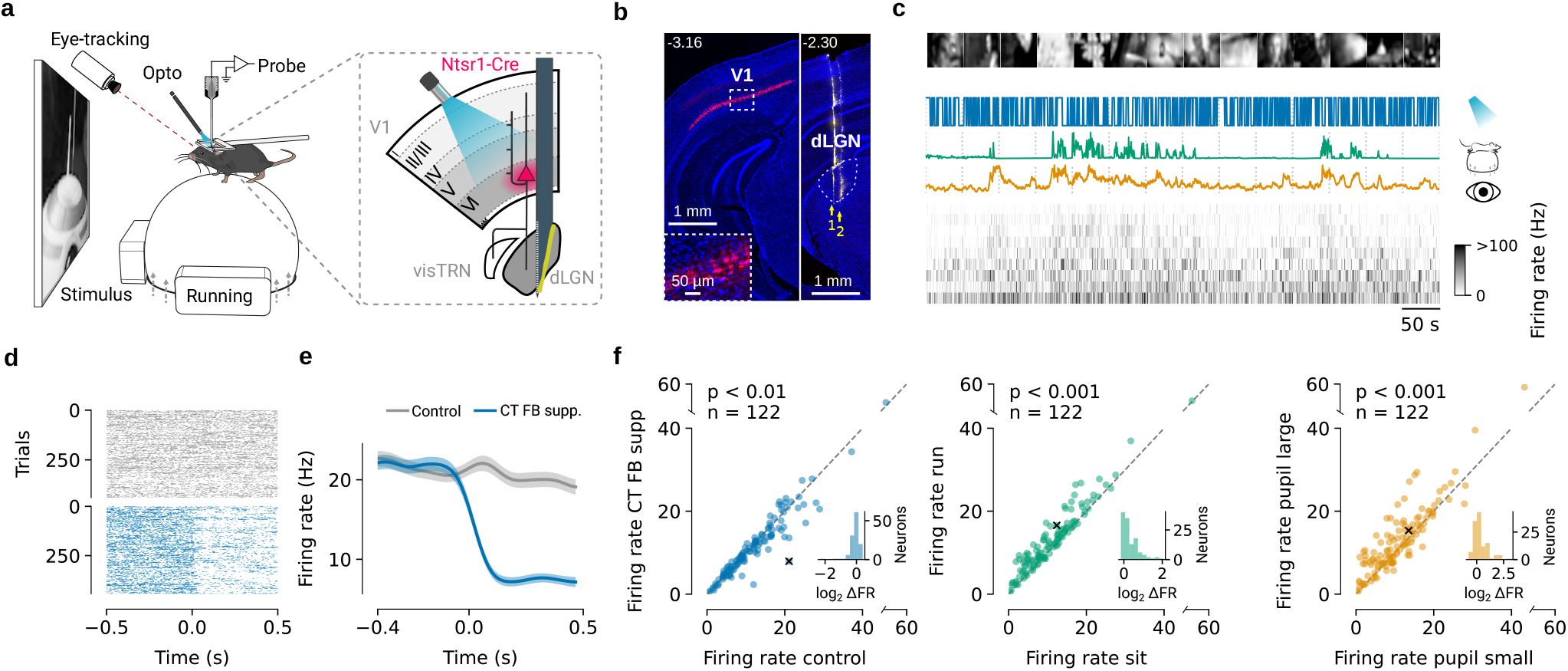
CT feedback and behaviour modulates dLGN responses to movies. (**a**) Schematic of the recording setup and photo-suppression of V1 L6 CT pyramidal neurons in Ntsr1-Cre mice with Cre-dependent AAV-stGtACR2-RFP. (**b**) Histology. *Left:* Coronal section near the V1 injection site, with stGtACR2-RFP expression (*red*) in Ntsr1+ somata. *Blue:* DAPI; scale bar: 1 mm. *Inset:* Magnification of area marked by dotted rectangle. Scale bar: 50 µm. *Right:* Coronal section of dLGN recording sites, with electrode tracks for two consecutive recording sessions (arrows 1 and 2) marked with DiI (*yellow*). Scale bar: 1 mm. *Dotted line*: dLGN contour. *Numbers on top*: position relative to Bregma in mm. (**c**) Snippet of an example dLGN recording. *Top to bottom*: example frames of the movie stimulus, photo-suppression pulse train (*blue*), running speed (*green*), pupil area (*yellow*), and time-varying firing rate (sorted by first principal component) of simultaneously recorded dLGN neurons. (**d**) Raster plot of responses of an example dLGN neuron, time locked to the onset of CT feedback photo-suppression (*blue*, OFF-ON transition) and to control periods without photosuppression (*grey*, OFF-OFF transition). Note that the example neuron illustrates the observed effect of CT feedback photo-suppression, but the size of the effect is not representative of that observed in the population of recorded dLGN neurons. (**e**) Corresponding PSTHs (*solid line*: average across trials, *shaded area*: standard error of the mean). (**f**) Effects of CT feedback photo-suppression (*left*), locomotion (*middle*), and pupil size (*right*) on dLGN mean firing rates.Example neuron from (d, e) marked with ×. *p* values denote results of a Wilcoxon signed-rank test, *n* = 122 neurons. *Insets:* Histogram of firing rate fold-change relative to control (Δ FR log_2_-ratio).

To develop initial insights into the modulations of dLGN responses by CT feedback and the other potentially modulating inputs, we aligned movie responses to onsets of photo-suppression, running, and pupil dilation. We found that certain neurons responded to the onset of CT feedback suppression with a substantial reduction in firing rate (**Figure 1d,e**, OFF-ON transitions in *blue*), while others showed milder effects or no modulation at all (**Figure 1f, left**). Despite the relatively small effect size, the overall reduction of dLGN firing rates during CT feedback suppression was genuine, as none of the recorded neurons in a control mouse without opsin expression showed systematic modulations at light onset (**Figure S1a, b**). Furthermore, neural responses aligned to time points in which the light was not switched on (i.e., OFF-OFF transitions in *grey*, **Figure 1d, e**) did not show any systematic modulation (**Figure S1c**). Finally, neurons with stronger CT feedback effects were closer to each other (**Figure S2**), as predicted by the topography of the corticothalamic system (*41, 7, 42, 43*). Consistent with previous findings (*11, 18, 10, 44*), we also found that during our movie stimulus, firing rates could gradually increase around transitions from sitting to running (**Figure S3a**), and during pupil dilation (**Figure S3b**).

These modulations of dLGN responses were also observed across the population of recorded neurons. Specifically, dLGN firing rates during the time of CT feedback suppression compared to periods without CT feedback suppression were reduced (*control vs. CT FB supp*. mean: 12.6 vs. 11.5 Hz, *p* = 1.24 × 10^−2^, paired Wilcoxon signed-rank test; **Figure 1f, left**), while time windows with running and dilated pupil were associated with an overall increase in average firing rates (*sit vs. run* mean: 11.0 vs. 12.8 Hz, *p* = 8.31 × 10^−16^, **Figure 1f, middle**; *small vs. large pupil* mean: 11.0 vs. 13.0 Hz, *p* = 4.98 × 10^−7^, paired Wilcoxon signed-rank test; **Figure 1f, right**). Although these modulations affected the recorded dLGN population on average, we also noticed considerable neuron-to-neuron variability. Indeed, for all three modulatory inputs, we found neurons that exhibited substantial variations in firing rates across the different conditions (**Figure 1f, insets**).

So far, it seems that our movie stimulus elicited various responses in dLGN, which were modulated by multiple additional inputs, this simple analysis on the mean firing rates does not take into account potential correlations between the different inputs. Consistent with previous studies that have shown that pupil size and running index partially overlapping behavioural states (*45, 46, 9*), we found a positive correlation between pupil diameter and running speed (*r* = 0.18 ± 0.17, mean ± SD; *p* < 0.001 for 7/10 experiments, permutation test; **Figure S3c**). Moreover, we found pupil size to also be influenced by stimulus brightness, where lower average intensity of movie frames was associated with larger pupil diameters (*r* = −0.41 ± 0.15, mean ± SD; *p* < 0.001 for 10/10 experiments, permutation test; **Figure S3c**). Pupil diameter, in turn, is known, to influence responses in the early visual system (*47, 48*).

### A spline-based GLM captures dLGN spatio-temporal RFs and their modulation by CT feedback and behaviour

To disentangle how the various potential influences shape the responses of dLGN neurons, we used a generalized linear model (GLM) (*27*) that predicted the neuron’s firing rate based on a combination of stimulus-driven and modulatory inputs (running speed, pupil size, and CT feedback suppression) (**Figure 2**). The model consisted of one linear kernel for each input, followed by a softplus function that accounted for response nonlinearities (**Figure 2a**). The shape of the stimulus kernel captured the neuron’s spatio-temporal RF, while the shapes of the modulatory kernels captured modulations of the neuron’s firing. We employed a GLM with a spline basis (*38*) in order to efficiently generate smooth kernels, rather than operating directly on the pixels of the visual stimulus or the discrete time bins of the additional inputs (**Figure S6a–d**). The GLM allowed us to effectively capture the temporal correlations between the inputs (see also above, **Figure S3c**), and the spatio-temporal correlations in pixel intensities in naturalistic stimuli (*49, 50*) (**Figure S4a**).

**Figure 2.**
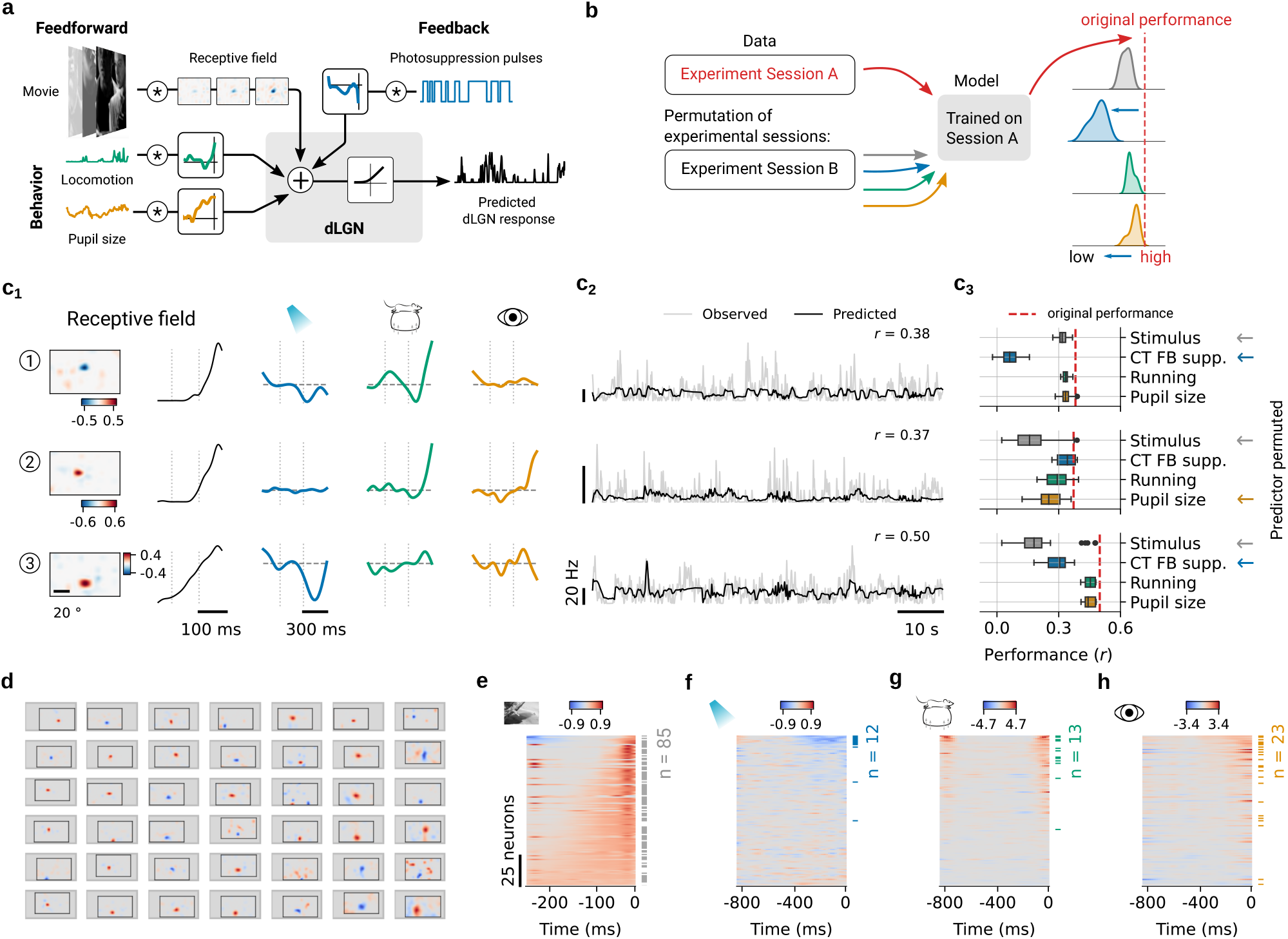
Spline-based GLM captured the RF, the influence of CT feedback and the impact of behaviour on responses of dLGN neurons. (**a**) Schematics of the spline-GLM model architecture. Firing rate in dLGN was predicted as a combination of kernel outputs summed at the linear stage and then passed through a softplus nonlinearity. Each modelled neuron had a kernel for the stimulus and kernels for the three modulatory inputs: run speed (*green*), pupil size (*orange*), and CT feedback suppression (*blue*). (**b**) Schematics of the permutation test (*53*) to evaluate the significance of the learned kernels. Model performance was evaluated by comparing the actual correlation (Pearson *r*) between predicted and observed firing rates to correlations when one of the inputs was taken from an unrelated experimental session (for movie, running, and pupil size) or randomly generated with the same statistics (for CT feedback suppression). (**c**) Three dLGN example neurons, their learned kernels, firing rate predictions, and outcomes of the permutation test (neuron 1 is the same example neuron as in **Figure 1d-e**). (c_1_) Spatial and temporal RF components separated by singular value decomposition (SVD, see Methods), along with kernels for the modulatory inputs. (c_2_) Observed (*gray*) versus predicted (*black*) firing rates during 80 s of movie presentation. (c_3_) Actual model performance (Pearson’s *r, red dashed line*) and performance for permuted stimulus (*gray*), CT feedback suppression (*blue*), running (*green*), and pupil size (*orange*) inputs. Kernels that contribute significantly to the model’s performance are marked with *←*. (**d**) Spatial RFs of example neurons with significant stimulus kernels. *Gray*: outline of common visual space (azimuth: −35–110 deg; elevation: −35–50 deg); *Solid lines*: monitor border. (**e**) Temporal RFs (SVD component of the stimulus kernel) in the recorded dLGN population, sorted by their area under the curve. (**f–h**) Modulatory kernels in the recorded dLGN population, sorted by their area under the curve, for CT feedback suppression (f), running (g), and pupil size (h). *Horizontal bars, side*: Neurons with significant kernels based on the permutation test. Panels (e–h) show data from all *n* = 122 neurons.

We trained and evaluated the spline-GLM on the recorded data set as follows: Given the known diversity of mouse dLGN feature selectivity (*51, 52*), we performed a separate hyperparameter search for each neuron. Hyperparameters included the number of spline bases for the stimulus and modulatory inputs and the weight for L1 regularisation (see Methods). All GLM fits in this study were cross-validated. The models analysed here were configured with the selected optimal hyperparameters and the reported performance is based on the held-out test set.

After model fitting, we assessed the predictive power of the learned kernel shapes using a session-based permutation test (**Figure 2b**). To keep the temporal statistics of the time series data intact, we provided the model with input data recorded on a different day (for the model inputs ‘stimulus’, ‘running’, ‘pupil size’) or with synthetic inputs generated with the same statistics as the original (for the ‘CT feedback suppression’ input). Subsequently, we compared for each input the actual model performance (Pearson’s *r*) against a distribution of model performances with that specific input permuted. Inputs were considered significant if the actual performance differed from the permuted performance with *p* ≤ 0.05.

We observed a rich diversity of learned GLM kernels for the dLGN neurons’ spatio-temporal RFs, the effects of CT feedback, and the behavioural variables. To begin with, we considered three example neurons (**Figure 2c**). Of these, the first had a negative spatial and a transient temporal stimulus response kernel, which contributed significantly to the model’s performance (*p* = 3.51 × 10^−2^, permutation test; **Figure 2c_1_**, **top**, same neuron as in **Figure 1d,e**). In addition, it had a significant negative kernel for CT feedback suppression (*p* = 2.67 × 10^−4^, permutation test; **Figure 2c_2_, c_3_**, **top**). In contrast, the running and pupil size kernels had minimal impact on this neuron’s model predictions (*p* > 0.05, permutation test). The second example neuron exhibited a positive spatial RF with an antagonistic surround (*p* = 2.31 × 10^−3^, permutation test; **Figure 2c_1_**, **middle**).While both behavioural kernels seemed to contribute (**Figure 2c_2_**, **middle**), only the kernel for pupil size reached significance in the permutation test (*p* = 1.13 × 10^−2^). This neuron was also not influenced by CT feedback suppression (*p* > 0.05; **Figure 2c_3_**, **middle**). Finally, example neuron 3 was primarily visually driven, with a positive RF centre and a more sustained temporal response kernel (*p* = 2.31 × 10^−3^, permutation test; **Figure 2c_1_**, **bottom**). It also had a significant negative kernel for CT feedback suppression (*p* = 2.67 × 10^−4^, permutation test; **Figure 2c_3_**, **bottom**). The other two modulatory kernels had only negligible influences (permutation test, *p* > 0.05; **Figure 2c_3_**, **bottom**).

The diversity observed in the three example neurons was also evident in the model fits across the population of recorded dLGN neurons, where we obtained a variety of spatio-temporal RFs, and combinations of modulatory influences. Assessing the learned spatio-temporal kernels, we found that ∼ 70% of the recorded dLGN neurons were visually responsive to our movie stimulus (85/122 neurons; permutation test visual stimulus, *p* ≤ 0.05; **Figure S6e**). For many of these neurons (representative examples in **Figure 2d**), the spline-GLM recovered spatial RF properties that were consistent with previous descriptions of mouse dLGN RFs obtained using artificial stimuli (*51, 54, 55*) and in many cases resembled those obtained from conventional sparse noise experiments (**Figure S4b,c**). These properties included various RF locations, RFs with either positive (66%, 56/85 neurons) or negative polarity kernels (34%, 29/85 neurons), a broad range of RF surround strengths (**Figure S4d,f**), and various RF centre sizes (**Figure S4e**). Furthermore, the GLM captured the well-known diversity of temporal response properties of dLGN neurons (*56, 51*), with some neurons showing more sustained, and others showing more transient temporal kernels (**Figure 2e**). Importantly, both the spatial and temporal GLM kernels matched well with the expected polarity and the dynamics obtained from clustering responses into sustained-OFF, Sustained-ON, and Transient groups during full-field luminance steps (**Figure S5**). This correspondence further underscores our model’s capacity to capture meaningful spatio-temporal RF properties and essential visual response characteristics of the recorded dLGN neurons.

We next assessed the learned kernels for the modulatory inputs. According to the permutation test, approximately 10% (12/122 neurons) of the recorded dLGN neurons were affected by CT feedback suppression, 11% (13/122 neurons) by running, and 19% (23/122 neurons) by pupil size (**Figure 2f–h, Figure S6f–h**). The overall direction of modulation for the significant kernels aligned with the modulation indices obtained directly from the data (**Figure 1f**): for the significantly modulated neurons, CT feedback suppression kernels were predominantly negative (**Figure 2f**), while running speed (**Figure 2g**) and pupil size (**Figure 2h**) kernels were predominantly positive. These results indicate that both CT feedback and behavioural state variables during natural movie viewing contribute to an overall increase of dLGN responses. Beyond the general sign of modulation, the learned model kernels also offered insights into the temporal dynamics of the modulatory influences: consistent with the well-known slow impact of pupil indexed arousal on responses in the visual system (*46, 45*), we found that kernels for pupil size had a more sustained profile compared to kernels for running modulations (time to half max kernel running vs. pupil size: 106.3 ms vs. 146.3 ms, Wilcoxon signed-rank test: *p* = 7.51 × 10^−3^, **Figure S6p**). This ability to learn differential kernels and a close correspondence between data-driven modulation indices and the impact of the modulatory inputs for model performance (**Figure S7d–g**) demonstrates that our GLM model was successful in extracting the impact of the various modulatory inputs.

Despite the spline-GLM’s overall success in predicting dLGN responses and learning biologically plausible kernels, it faced challenges in capturing fast modulations and response peaks, as typical also for traditional GLMs (*26, 57*). We also observed increased variance at the left side of some kernels particularly evident for running modulated neurons (**Figure 2e–h** and **Figure S6i–l**). This could potentially stem from typical boundary artefacts associated with splines (**Figure S6a–d**). Nevertheless, despite these challenges, incorporating running information proved beneficial, as confirmed by the permutation test (**Figure S6g**). Reassuringly, these running modulated neurons showed reduced response reliability in repeated stimulus trials (**Figure S6m,n**) as their activity was substantially influenced by locomotion state. In line with a previous study (*58*), these neurons also had overall low firing rates (**Figure S6o**).

Is the contribution of modulatory inputs to dLGN neurons’ activity consistent and strong enough to improve the prediction of dLGN responses? While dLGN has long been known to exhibit state-dependent changes in firing (*15, 11, 59, 18*) and to receive extensive feedback from cortex (*60, 61*), the impact of these influences, in particular during viewing of naturalistic stimuli, is not well understood. Thus, to quantitatively assess the contributions of CT feedback suppression and behavioural variables, we compared the performance between our model including all inputs and reduced variants of the model with only a subset of inputs (**Figure 3a,b**). Starting with the ‘Stimulus only’ model that only considered the stimulus as input, we found that incorporating one or more modulatory inputs increased the correlation between observed and predicted dLGN responses (ANOVA: *p* = 0.022; **Figure 3a**). In particular, adding pupil size or a combination of two or more predictors showed a significantly better performance than the ‘Stimulus only’ model (‘Stimulus only’ vs. ‘Stimulus + Pupil size’: 0.186 vs. 0.235, Wilcoxon signed-rank test: Bonferroni corrected *p* = 3.39 × 10^−6^; ‘Stimulus only’ vs. ‘Full model’: 0.186 vs. 0.249, Wilcoxon signed-rank test: Bonferroni corrected *p* = 2.08 × 10^−7^; **Figure 3a**). Note that even in the full model we still observed neurons with suboptimal predictions, maybe due to low response reliability (**Figure S6m,n**) and sparse firing (**Figure S6o**).

**Figure 3.**
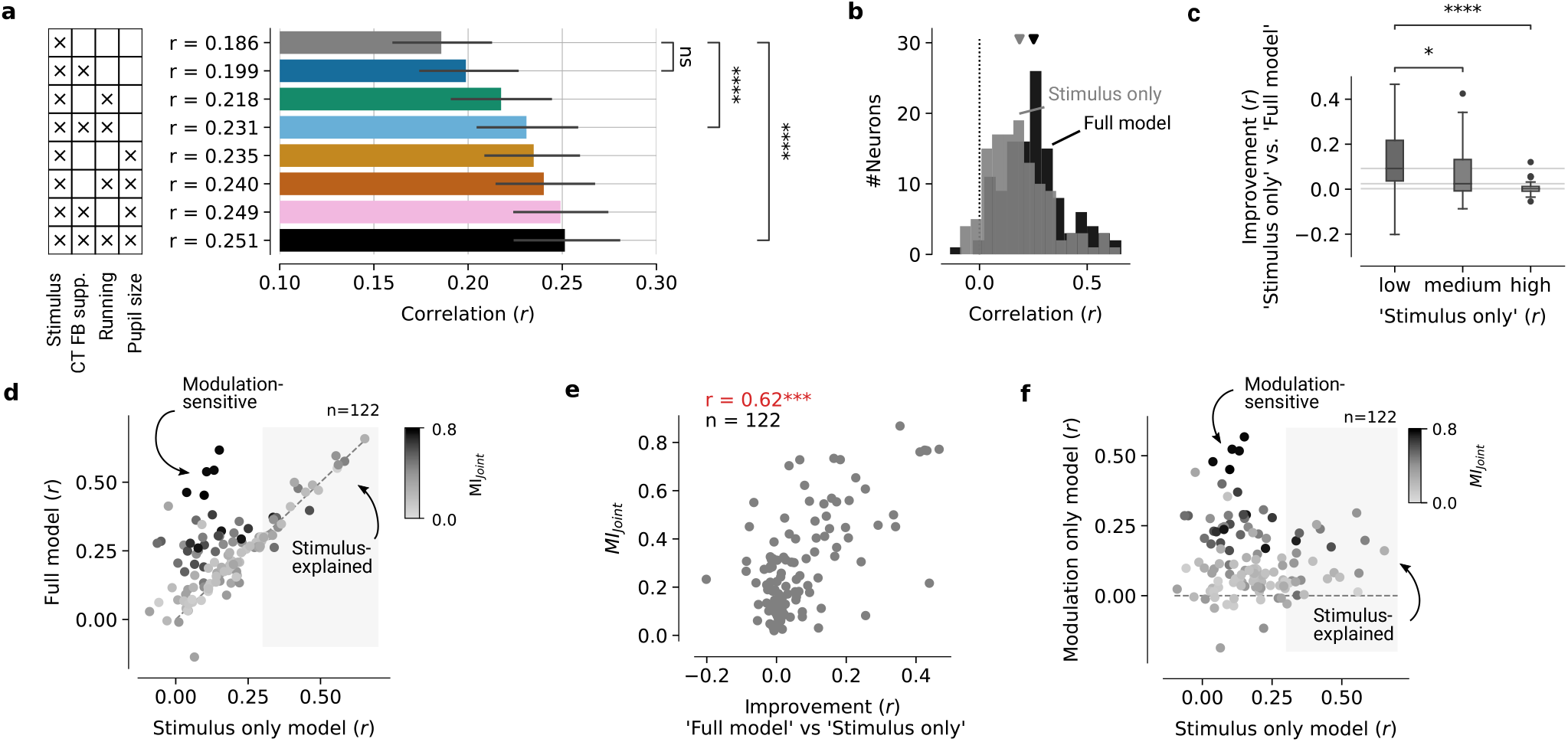
Adding model predictors for modulatory inputs improves the performance for a subgroup of poorly visually responsive dLGN neurons. (**a**) Comparison of model performance for models with different sets of inputs indicated in the table on the left, sorted by performance. Bonferroni corrected *p*-values of paired Wilcoxon signed-rank test: **** *p* ≤ 1.0 × 10^−4^; ‘ns’ non-significant. Error bars: 95% confidence intervals. (**b**) Comparison of model performance on the test set (Pearson’s *r*) for the ‘Full model’ (*black*, inputs: stimulus, CT feedback suppression, run speed, pupil size) and the ‘Stimulus only’ model (*gray*). *Arrow heads*: mean performance (*n* = 122 neurons). (**c**) Improvement in model performance observed with adding predictors for modulatory inputs (‘Full model’−’Stimulus only’) for neurons grouped by the (low/medium/high) performance of the ‘Stimulus only’ model. Bonferroni corrected *p*-values of paired Wilcoxon test: *\*p* ≤ 0.05; * * * * *p* ≤ 1.0 × 10^−^4. (**d**) Comparison of model performances on the test set for all neurons in the ‘Stimulus only’ model and the ‘Full model’. Darker colours indicate stronger joint modulation by CT feedback, running, and pupil size (*MI*_*Joint*_) estimated directly from the data without model (see Methods). Arrows indicate the group of neurons that benefits from adding CT feedback, running, and pupil size (‘Modulation-explained’) and the group that does not improve (‘Stimulus-explained’). (**e**) Improvement in model performance observed with adding predictors for modulatory inputs (‘Full model’−’Stimulus only’) and the joint modulation by CT feedback, running, and pupil size estimated directly from the data without model (see Methods). (**f**) Performances of a model that considers only the stimulus as input (‘Stimulus only’) and a model without the stimulus (‘Modulation only’). Arrows indicate ‘Modulation-explained’ and ‘Stimulus-explained’ neurons. Darker colours indicate stronger joint modulation by CT feedback, running, and pupil size estimated directly from the data without model. Panels (d–f) show data from all *n* = 122 neurons.

Indeed, when we compared the prediction performance of the ‘Stimulus-only’ model with the ‘Full model’ (**Figure 3b**), we noticed that the inclusion of the modulatory inputs did not merely shift the distribution to higher performances. Instead, a closer examination allowed us to identify a subgroup of neurons whose responses could be predicted only poorly by the visual stimulus, but which showed substantial improvements in their response prediction with the inclusion of modulatory factors (**Figure 3c,d**; we call them the ‘Modulation-sensitive’ group; **Figure S7a–f**). In contrast, another subgroup showed no improvement when we added the modulatory inputs (**Figure 3c,d**; ‘Stimulus-explained’ group). To test the extent to which this reflected a ceiling effect in explainable variance, we fitted models that had only the modulatory factor as inputs but not the stimulus (‘CT FB supp.’ + ‘Running’ + ‘Pupil size’; ‘Modulation only’ models; **Figure 3f**). For these models, neurons substantially affected by modulatory inputs still performed well (**Figure 3f**, ‘Modulation-sensitive’, dark dots), whereas those well-explained by the visual stimulus exhibited lower performance, although not zero (**Figure 3f**, ‘Stimulus-explained’, light grey dots). This indicates that even in the ‘Stimulus-explained’ group, modulatory inputs might have some, albeit generally weak effect that was obscured in the full model; alternatively, some modulatory factors might contain stimulus information, in particular pupil size, which is known to be related not only to arousal, but also to stimulus brightness (**Figure S3c**). Taken together, our analysis of model performance suggests that dLGN neurons are explained by the visual stimulus, behaviour and CT feedback, albeit with considerable heterogeneity.

### CT feedback is enhanced in the absence of a patterned visual stimulus

Our model showed only a relatively small subset of dLGN neurons with significant effects of CT feedback. Could the reason for this be that the responses elicited by the rich naturalistic movie stimulus might have dominated dLGN activity relative to the effects of CT feedback? We thus predicted that CT feedback effects might be stronger without patterned stimulus input (**Figure 4a**).

**Figure 4.**
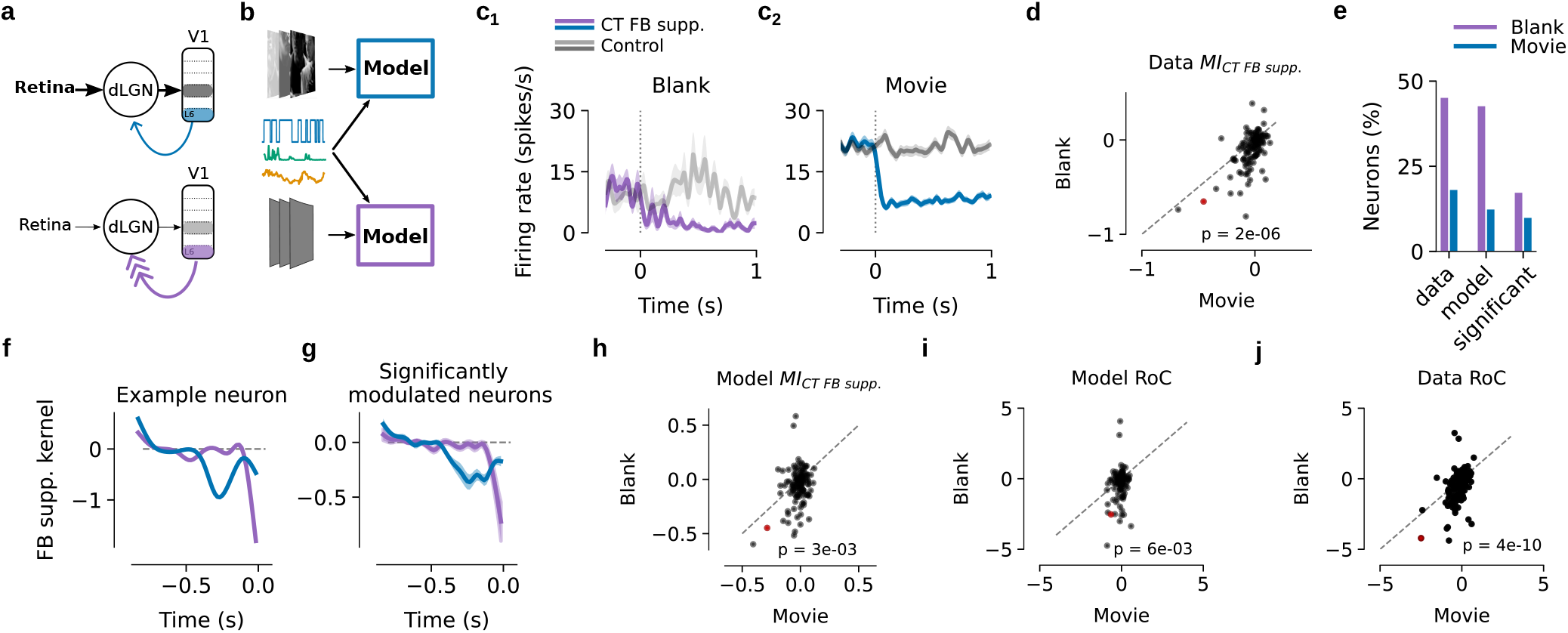
The effect of CT feedback is dependent on the presence or absence of the visual stimulus. (**a**) Schematics of the hypothesis that the absence of a patterned visual stimulus elicits weaker stimulus-driven input to dLGN, which is in turn accompanied by stronger CT feedback. Note that the schematics should not imply stronger L6 CT pyramidal neuron firing, but that the net effect of CT feedback on dLGN firing is stronger in the absence of a patterned visual stimulus. (**b**) Schematics of fitting the spline-GLM model separately during movies (*top*) vs. blank periods (*bottom*). (**c**) Effects of CT feedback suppression. (c_1_) Mean PSTHs time locked to onset of photosuppression for one example dLGN neuron (same example neuron as in **Figure 1d-e**) during blank periods (gray screen) flanking the movie presentation. The shaded area represents the SEM. (c_2_) Same as (c_1_), during movie presentation. *Purple, blue*: PSTH during CT feedback suppression (OFF-ON transition), *light grey, dark grey*: PSTH during control condition (OFF-OFF transition). (**d**) Comparison of *MI*_*CT FB supp*._ during blanks vs. movies for the recorded dLGN population (number of neurons *n* = 122). (**e**) Percentage of recorded dLGN neurons modulated by CT feedback during the two stimulus conditions. Three modulation metrics were separately considered to count the modulated neurons. A neuron was considered modulated (1) based on data: |*MI*_*CT FB supp*._| from (d) ≥ 0.1, (2) based on model predictions: |*MI*_*CT FB supp*._| from (h) ≥ 0.1, or (3) based on model performance: permutation test *p* ≤ 0.05. Notably, all three modulation metrics consistently revealed a higher proportion of neurons displaying CT feedback modulation during the blank condition compared to the movie condition. (**f**) CT feedback suppression kernel for the example dLGN neuron in (c_1_). The model was either trained on the data from movie presentation (*dark blue*) or blank periods (*light blue*). (**g**) Same for all significantly CT feedback modulated neurons (*n*_movie_ = 12, *n*_blank_ = 21). Solid lines represent the mean of the kernels and transparent surrounds represent the standard error of the mean (SEM). (**h**) Comparison of *MI*_*CT FB supp*._ for blanks vs. movies calculated from simulated data using the fitted model for the dLGN population to the two stimuli. (**i**) Comparison of the Rate of Change (RoC) of model-predicted neurons’ responses to CT feedback suppression during movie presentation vs. blank periods. (**j**) Same as (i), for the recorded data. Panels (h–j) show data from all 122 neurons.

To test this prediction, we expanded our analyses to the period of blank screen stimulation flanking the movie presentation, and compared the effects of CT feedback suppression in the recorded data and in models fit separately to responses during movies vs. blank periods (**Figure 4b**). Consistent with our prediction, we indeed observed that CT feedback suppression reduced the firing rates of individual example neurons more strongly during blanks (*MI*_*CT FB supp*._: −0.65, **Figure 4c_1_**) compared to movies (*MI*_*CT FB supp*._: −0.45, **Figure 4c_2_**). This observation held true across the recorded dLGN population (mean *MI*_*CT FB supp*._ −0.09 vs. −0.03; *p* = 2.5 × 10^−6^, Wilcoxon signed-rank test; **Figure 4d**). Furthermore, during blank periods, more dLGN neurons were affected by the suppression of CT feedback (45% vs. 18% with |*MI*_*CT FB supp*._| ≥ 0.1; **Figure 4j**). The stronger effect of CT feedback suppression could not be explained by the difference in overall firing rate between the two stimulus conditions (**Figure S8a-b**), nor was it related to the different number of optogenetic pulses (**Figure S8c**).

Is the stronger modulation by CT feedback suppression during blank periods vs. movies also captured by our spline-GLM model? We indeed found that the model learned a more negative CT feedback suppression kernel for the blank condition compared to the movie, both in the example neuron (peak amplitude −1.8 vs. −0.94; **Figure 4f**) as well as in the population of neurons with a significant CT feedback suppression kernel (−0.86 vs. −0.46, *p* = 0.044, Mann-Whitney-U test, **Figure 4g**; all neurons: −0.4 vs. −0.2, *p* = 5 × 10^−8^, Wilcoxon signed-rank test). To avoid potential confounds of firing rates on kernel amplitudes (**Figure S8d,e**) and enable a more direct comparison to the recorded data, we used the model predictions in the two stimulus conditions and calculated model-derived *MI*_*CT FB supp*._, analogously to those based on the recorded data. Consistent with our data-driven observations, we found that CT feedback suppression reduced modelled responses more strongly during the blank periods (modelled *MI*_*CT FB supp*._, −0.057) compared to movies (−0.017; *p* = 0.002, Wilcoxon signed-rank test, **Figure 4h**)

Inspecting the learned model kernels (**Figure 4f,g**) suggested that, besides amplitude, the dynamics of the CT feedback suppression effects might also differ between visual stimulus conditions. Specifically, the spline-GLM learned a kernel characterised by a faster and briefer time course for blank periods compared to movie stimulation. This observation could be quantified by calculating the rate of change (RoC) of the neurons’ predicted and actual responses after photosuppression (see Methods). Indeed, we found that the effect of CT feedback suppression was faster during blank periods (model-based RoC: *p* = 0.004, Wilcoxon signed-rank test, **Figure 4i**; data-based: *p* = 9 × 10^−12^, Wilcoxon signed-rank test,**Figure 4j**). In conclusion, the fitted spline-GLM models revealed that the observed impact of CT feedback suppression was weaker and slower during the presence of a rich patterned visual stimulus in comparison to blank periods, suggesting that the observed effect of CT feedback depends on the visual stimulus characteristics.

### CT feedback affects stimulus decoding

Finally, we sought to better understand if and how the suppression of CT feedback affected how dLGN encoded visual information (**Figure 5**). Specifically, we tested the degree to which suppression of CT feedback might affect the decoding of movie information based on the responses of single dLGN neurons, compared to the control condition where feedback was intact.

**Figure 5.**
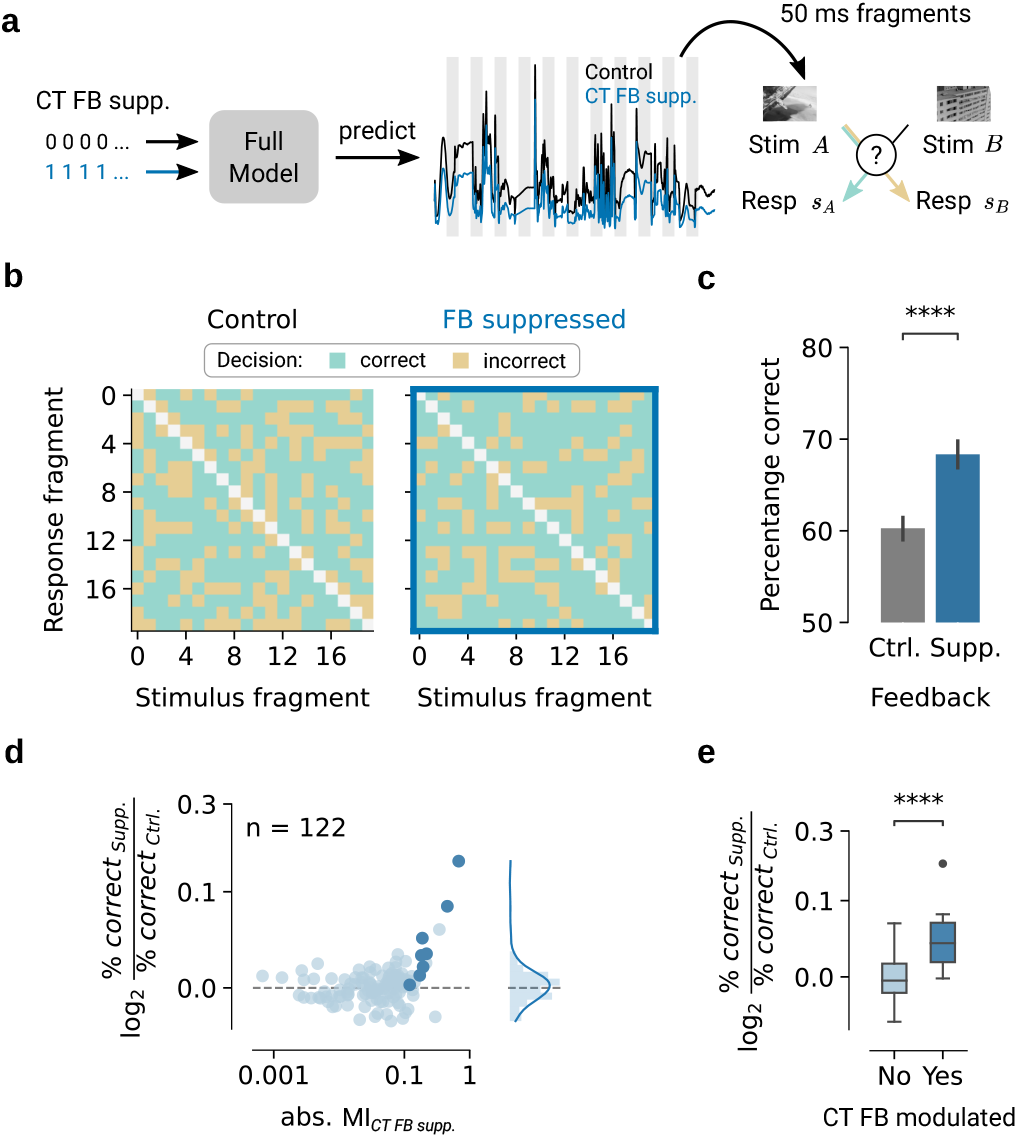
Two-alternative-forced-choice decoder shows better stimulus discrimination for CT feedback modulated neurons when feedback is suppressed. (**a**) Schematics of the decoder: the trained model was used to simulate responses with CT feedback being either on (*blue*) or off (*black*; 70 s example trace). Random 50 ms stimulus fragments (*A* and *B*) were chosen, each with corresponding simulated responses. The decoder’s task was to determine which stimulus was more likely based on the observed responses. (**b**) Decision matrix for one example neuron with 20 random stimulus fragments and their simulated responses. *Green*: correct pairing of stimulus and response; *yellow*: incorrect pairing. *Left*: control condition; *right*: CT feedback suppressed condition. (**c**) Percentage of correct decisions in the control and the CT feedback suppressed condition, same example modelled neuron as in (b). Error bars indicate 95% confidence intervals. (**d**) Decoder performance across all neurons (*n* = 122) as a function of the amount of modulation by photosuppression observed in the experiment. *Dark blue*: neurons with a significant CT feedback kernel. (**e**) Relative decoder performance during CT feedback suppression and control conditions, split according to whether neurons were significantly modulated by CT feedback suppression (*dark blue*) or not modulated (*light blue*).

To address our question, we used the trained model (‘Full model’) to describe how individual dLGN neurons encode stimulus information (**Figure 5a**). In this way, we were able to isolate the effects of CT feedback suppression *in silico*, by keeping stimulus, running and pupil inputs intact, and only changing CT feedback suppression (setting the CT feedback suppression either OFF (0) or ON (1)). We then performed a decoding experiment, using simulated model responses to discriminate between video clips in a two-alternative forced choice (2AFC) setting (*30*). We randomly selected 50 ms fragments of stimulus-response-pairs from the test set that had not been used for model fitting. To discriminate between two movie clips, we used the maximum-likelihood decoding rule from Pillow et al. (2005) (*30*). The decision of the 2AFC decoder was directly derived from the model likelihood for the correct and incorrect pairing (see Methods and (*30*)). We considered the 2AFC decoder to make a correct decision if the likelihood for the correct pairing was higher compared to the incorrect one. We found that the decoder made more correct decisions in the FB suppressed condition compared to the control condition (**Figure 5b**, 20 representative stimulus-response pairs for one example neuron; **Figure 5c**, across all 100 stimulus pairs 68.34 % vs. 60.27 %, *p* = 1.20 × 10^−12^, Wilcoxon rank sum test).

To quantify the discrimination performance for each neuron, we computed the log ratio of the percent of correct choices in the FB suppressed condition and the control condition (**Figure 5d**), with positive log ratios indicating better decoding performance in the CT feedback suppressed state. The observed log-ratios near zero for the majority of neurons suggest that there was no difference in stimulus information between control conditions and CT feedback suppression (average 1.003±0.021, mean±SD; **Figure 5d**). However, we found significantly higher log-ratios in neurons modulated by CT feedback compared to non-modulated neurons (*p* = 5.84 × 10^−14^, skew test against null; **Figure 5d, right** and **Figure 5e**). Consequently, our model predicts that with intact CT feedback, short duration activity of individual dLGN neurons can be less informative about the visual stimulus.

## Discussion

Here, we quantitatively characterised how dLGN responses during viewing of a naturalistic movie are influenced by the combination of visual stimulus-related inputs, CT feedback and behavioural state. We modelled the responses of individual dLGN neurons using a data-efficient GLM, which predicted the spatio-temporal RFs and response modulations by CT FB suppression, the animal’s run speed, and pupil size. We found that overall model performance improved when including the modulatory predictors, in particular for a subpopulation of dLGN neurons whose responses were poorly explained by the visual stimulus. Guided by the model, we found that the effect of CT FB suppression depended on stimulus type, being relatively stronger, more prevalent, and faster in the absence of a patterned visual stimulus. Finally, using Bayesian decoding, we found that the responses of a subset of CT-feedback-sensitive dLGN neurons contained more visual information when CT feedback was suppressed. Together, our results show that the activity of dLGN neurons is governed by a combination of stimulus-driven and modulatory inputs and is mediated by their visual responsiveness and the stimulus type.

### Using a spline-GLM for modelling thalamic responses

Our work extends previous approaches that modelled thalamic processing of naturalistic visual stimuli (e.g., (*62, 37, 36*)). In particular, our framework allowed us to test model variants with different combination of predictors, which revealed a subpopulation of dLGN neurons whose responses were poorly predicted by the movie stimulus and more strongly affected by the modulatory inputs. These might correspond to a previously reported set of neurons in mouse dLGN, amounting to 30–40%, with poor or unclear visual feature selectivity (*63, 56*). Given that this subpopulation also contained an over-representation of low-firing neurons, their relatively stronger modulation (see also (*58*)) might be indicative of a tight inhibitory / excitatory coupling in the recorded network (*64*). Such tight coupling in the thalamo-cortico-thalamic network might be mechanistically achieved through joint modulations of dLGN relay cells, and inhibitory neurons in dLGN and the thalamic reticular nucleus by CT feedback and neuromodulation (*1, 4, 7, 9*).

The general success of our model is indicated by the close neuron-by-neuron correspondence between RF properties and response types derived from the model and more traditional analyses and stimuli, and by the prominent relationship between learned model kernels and data-driven modulation indices for CT feedback suppression, run speed and pupil size. Yet, while our data-driven modulation indices generally matched well with the magnitudes of modulatory influences previously observed in mouse dLGN (*11, 18, 10, 7, 14*), we were surprised to find only a relatively small fraction of dLGN neurons with significant kernels for CT feedback suppression, running and pupil size. Reasons for the small fraction might be at least threefold: (1) the conservative method for assessing significance in the model derived from session-based permutation tests (*53*), (2) the continuous nature of inputs to the model rather than a split into extreme conditions for some of the modulation indices (e.g., locomotion slower than 0.25 cm/s was considered as sitting but only faster than 1 cm/s was considered as running (*11, 18*)), (3) the direct suppression of L6 CT feedback through the light-gated chloride channel stGtACR2 (*39*), which might have yielded comparatively weaker effects (see also (*10, 65*)) than alternative approaches recruiting powerful intracortical inhibition through photostimulation of inhibitory V1 neurons (*7, 6, 66, 10, 65, 9*).

In the future, our spline-GLM could be extended by thalamic mechanisms, such as the fast adaptation of integration time according to luminance and contrast (*67, 62*), accounting for the constant changes in spatial and temporal integration elicited by dynamic natural stimuli (*68*). Further, by combining the model inputs, one could test for interactions, for instance to clarify the dependence of CT feedback effects and behavioural state-related modulations as proposed by some studies (*65*). Incorporating adaptive amplitudes of input and post-spike kernels (*69*) could differentiate tonic and burst spiking behaviours, characteristic of thalamic neurons (*70*) and known to be affected by both stimulus-related (*71*) and modulatory inputs (*72, 15, 11, 10*). Finally, alternative modelling approaches based on deep neural networks (DNNs) (*73, 74, 75*) promise to better capture non-linear dynamics and provide faster computations, potentially enhancing our capabilities to emulate neuronal responses and complex neural interactions, yet likely at the expense of interpretability and uncontrolled biases (*76, 77*). Some recent efforts have been made to increase the interpretability and biological plausibility of such models (*78, 79*), at times, by crafting an integrative model combining the advantageous non-linearity of DNNs with linear models like ours (*80, 81*).

### Spatio-temporal RFs and modulatory inputs of mouse dLGN neurons

Through a combination of analysing receptive fields obtained from the model using a naturalistic movie stimulus, a noise stimulus, and more traditional RF mapping techniques, we identified both expected and unexpected types of dLGN spatio-temporal RFs. Indeed, consistent with previous studies quantifying RFs in mouse dLGN to simple stimuli (*51, 56, 55, 54*), our model learned circular spatial RFs for the majority of neurons, which often consisted of a single domain resembling the well known ON and OFF fields. Reminiscent of a study showing that retinal ganglion cells can reverse the polarity of their RFs in response to different natural images (*82*), we found that some dLGN neurons had an opposite RF polarity when characterised with simple luminance steps or the movie stimulus. Finally, some of the RFs obtained with our modelling approach had a complex spatial structure, and might thus correspond to a subset of dLGN neurons that had been previously noted to lack clearly localised RFs (*56, 54*). Future studies, for instanced based on the “maximally exciting image” approach initially applied to mouse V1 (*83*), are needed to verify to which degree these complex RF structures indeed reflect complex feature selectivity of dLGN neurons or might be potential consequences of the modelling approach.

Despite its frequent portrayal as a relay station, the dLGN of the thalamus has long been recognised as one of the earliest stages in the visual system that integrates visual representations with additional information (*3, 15, 12, 4, 84*). Corroborating this, we found that the performance of our model generally improved with the inclusion of modulatory inputs, albeit with considerable neuron-by-neuron diversity. Previous studies have already reported differential effects of behavioural state and arousal, and could relate them to the neurons’ feature selectivity (*85, 18, 19*). In dLGN, the firing of neurons with nonlinear responses to high spatial frequencies and with transient ON responses seem preferentially enhanced by locomotion (*18*). In addition, retinal boutons preferring low spatial frequencies and luminance decrements seem preferentially suppressed by arousal (*19*). The neuron-by-neuron diversity in the impact of the modulatory inputs might thus serve to enhance particular visual inputs during active states.

### CT feedback effects on single dLGN neuron stimulus encoding and decoding

Our observation of faster and relatively stronger CT feedback effects during blank periods compared to movie viewing contributes to the growing appreciation that CT feedback effects on dLGN firing rates seem to be stimulus-dependent, and potentially overridden by strong visual stimulation (see also (*86, 10*)). Indeed, the effect of CT feedback seems to be most potent in the absence of patterned stimulus input, both in mice (this study and (*41, 10*) for related findings with gratings) and ferrets (*86*), which might point to a common mechanism across species. We propose that a greater influence of CT feedback suppression during spontaneous activity than movie viewing could arise from a differential engagement of direct excitatory and inhibitory feedback pathways during these visual stimulus types. Supplying thalamic relay neurons with different ratios of excitatory and inhibitory conductances, mimicking the impact of modulatory inputs, can shift their input/output function, such that the same somatic input can generate markedly different spiking responses (*87, 88*). Future studies will need to use more subtle manipulations of stimulus type, including contrast and spatial structure, together with pathway-specific CT feedback suppression, to test the hypothesis that CT feedback might be most effective under challenging sensory conditions. In addition, future studies will profit from simultaneous recordings of V1 and dLGN to further disentangle the origin and potential interaction of CT feedback and other modulatory influences.

We also observed that suppressing CT feedback enhanced the decoding of visual information in individual, feedback-sensitive neurons. This was surprising given prior research associating cortico-cortical feedback with attention and enhancements of stimulus encoding (e.g., (*89, 90, 91, 92, 93*)). What could be potential reasons why our model predicted improved decoding during CT feedback suppression? Past research has proposed that one role of CT feedback could be to linearise the input-output relationship of thalamic neurons through the injection of synaptic noise (*87, 88*). This linearisation would enhance the excitability of thalamic neurons to weaker inputs, but would additionally push them into more unreliable firing regimes. This could be one potential explanation why our decoder operating on single stimulus segments performed worse with CT feedback intact. A shift towards more unreliable firing regimes by CT feedback would also be in line with results of a previous study (*10*), showing that CT feedback during repeated movie segments increased the trial-by-trial variability of dLGN neurons’ responses. Such increased variability in individual dLGN neurons could be offset and even exploited by the strong convergence in the thalamocortical system (*94, 95*). Specifically, synaptic noise mediated by CT feedback might allow to extract from the pooled afferent signal stimulus-related information with better resolution and with enhanced sensitivity to weaker inputs (*87, 88*). Thus, in the future, a more realistic decoder would consider local populations of thalamocortical neurons, to test the hypothesis that, through CT feedback mediated synaptic noise, cortex regulates the trade-off between heightened sensitivity towards ambiguous inputs and reliable representation of clear inputs.

In conclusion, our results add to the growing body of evidence that dLGN activity is influenced not only by visual inputs but also by modulatory influences from CT feedback and behavioural state. Our work presents an important step towards a quantitative understanding of how dLGN responses to complex, naturalistic stimuli are shaped by the simultaneous influences of stimulus-related feedforward inputs, CT feedback and behaviour.

## Acknowledgements

This research was supported by the Deutsche Forschungsgesellschaft (DFG) Sonderforschungsbereich (SFB) 1233, *Robust Vision: Inference Principles and Neural Mechanisms*, Teilprojekt (TP) 13, project number: 276693517 (L.B., P.B.), by SPP2041 (BU 1808/6-1 and BU 1808/6-2; L.B.), the RTG 2175 “Perception in context and its neural basis” (L.B.) and the Hertie Foundation (P.B.). Lisa Schmors was supported by the International Max Planck Research School for Intelligent Systems (IMPRS-IS). We thank E. Froudarakis (A. Tolias Lab, Baylor College of Medicine, Houston, TX) for providing of the movie stimulus files, S. Renner for helping with the database implementation, and A. Ecker and T. Euler for discussion of the stimulus design. Thanks also go to M. Sotgia for lab management and support with animal handling and histology, S. Schörnich for IT support, and B. Grothe for providing excellent research infrastructure.

## Author contributions

Conceptualization, L.B., P.B., S.S.; Methodology, L.S., Y.B., A.K., Z.H., D.C. P.B., S.S., L.B.; Software, L.S., A.K., Y.B., Z.H., D.C.; Formal Analysis, L.S., A.K., Y.B.; Investigation, Y.B., A.K., L.M.; Resources, L.B., P.B.; Data Curation, L.S., A.K., Y.B., L.M., D.C.; Writing – Original Draft, L.B., L.S., A.K., Y.B.; Writing – Review & Editing, all authors; Visualization, L.S., A.K., Y.B.; Supervision, L.B., P.B., S.S.; Project Administration, L.B., P.B.; Funding Acquisition, L.B., P.B.

## Data and code availability

Data and code will be made available upon submission.

## Methods

All procedures complied with the European Communities Council Directive 2010/63/EU and the German Law for Protection of Animals, and were approved by local authorities, following appropriate ethics review.

### Surgical procedures

Experiments were carried out under under Licence ROB-55.2-2532.Vet_02-17-40 in 4 adult Ntsr1-Cre mice (median age:15.5 ± 6.45 weeks; B6.FVB(Cg)-Tg(Ntsr1-cre)GN220Gsat/Mmcd; MMRRC) of either sex. Stereotactic surgeries were performed to implant a head-post for head-fixation, implant a ground/reference screw for electrophysiology, inject a virus for optogenetic feedback manipulation, and drill a craniotomy for acute electrode insertions.

### Stereotactic surgery preparation and initiation

Thirty minutes prior to the surgical procedure, mice were injected with an analgesic (Metamizole, 200 mg/kg, sc, MSD Animal Health, Brussels, Belgium). To induce anesthesia, animals were placed in an induction chamber and exposed to isoflurane (5% in oxygen, CP-Pharma, Burgdorf, Germany). After induction of anesthesia, mice were fixated in a stereotaxic frame (Drill & Microinjection Robot, Neurostar, Tuebingen, Germany) and the isoflurane level was lowered (0.5 %–2 % in oxygen), such that a stable level of anesthesia could be achieved as judged by the absence of an interstitial reflex. Throughout the procedure, the eyes were covered with an eye ointment (Bepanthen, Bayer, Leverkusen, Germany) and a closed loop temperature control system (ATC 1000, WPI Germany, Berlin, Germany) ensured that the animal’s body temperature was maintained at 37° C. At the beginning of the surgical procedure, an additional analgesic was administered (Buprenorphine, 0.1 mg/kg, sc, Bayer, Leverkusen, Germany) and the animal’s head was shaved and thoroughly disinfected using iodine solution (Braun, Melsungen, Germany). Before performing a scalp incision along the midline, a local analgesic was delivered (Lidocaine hydrochloride, sc, bela-pharm, Vechta, Germany). The skin covering the skull was partially removed and cleaned from tissue residues with a drop of H_2_O_2_ (3 %, AppliChem, Darmstadt, Germany). Using four reference points (bregma, lambda, and two points 2 mm to the left and to the right of the midline respectively), the animal’s head was positioned into a skull-flat configuration for the further steps.

### Virus injection

In order to suppress V1 L6 CT FB selectively and reversibly, we conditionally expressed the chloride-conducting channelrhodopsin stGtACR2 (*39, 96, 97*) in L6a CT pyramidal cells (*84, 60, 61*) by injecting AAV-stGtACR2-RFP into the left hemisphere V1 of Ntsr1-Cre mice (*40, 98, 99*) (**Figure 1a**). Ntsr1+ neurons are known to correspond with > 90% specificity to L6 CT pyramidal cells (*6, 100, 101*). Furthermore, the opsin *stGtACR2* restricts expression to somata and the axon-initial segment which prevents possible accidental axonal depolarization due to a differential Cl^-^ ion reversal potential across different neuronal compartments (*102, 103, 104*). It also offers improved photocurrents and higher sensitivity, which are of particular relevance to manipulating deeply located L6 CT neurons, while avoiding light artifacts and tissue damage arising from excessive light intensities (*104*).

Before surgery, the Cre-dependent, stGtACR2-expressing *adeno-associated virus* (AAV) vector (pAAV_hSyn1-SIO-stGtACR2-FusionRed, Addgene, #105677) stock solution was diluted to 5×10^11^ gc/ml titers, and aliquotted to 4 *µ*L.

During surgery, aliquots were front-loaded into a glass pipette mounted on a Hamilton syringe (SYR 10 *µ*L 1701 RN no NDL, Hamilton, Bonaduz, Switzerland), controlled by the Injection Robot of the Neurostar Stereotax. After performing a small craniotomy for injection (100 *µ*m diameter), we injected 300 nl of virus solution into V1 (2×50 nl shots injected at a rate of 50 nl / 30 s at a respective depth of 900 *µ*m, 800 *µ*m and 700 *µ*m below the brain surface.

### Head-post and ground and reference screw implantation

For implant fixation, the exposed skull was covered with OptiBond FL primer and adhesive (Kerr Dental, Rastatt, Germany) omitting three locations: V1 (AP: −3.28 mm, ML: −2.4 mm), dLGN (AP: −2.3 mm, ML: −2 mm), and a position roughly 1.5 mm anterior and 1 mm to the right of bregma, designated for a miniature ground and reference screw.

A custom-made lightweight stainless steel head bar was positioned over the posterior part of the skull such that the round opening in the bar was centered on V1/dLGN. The head bar was attached with dental cement (Ivoclar Vivadent, Ellwangen, Germany) to the primer/adhesive. The opening was later filled with the silicone elastomer sealant Kwik-Cast (WPI Germany, Berlin, Germany). Then the miniature screw (00-96 × 1/16 stainless steel screws, Bilaney), which served both as ground and reference that was soldered to a custom-made connector pin, was implanted.

### Post-surgical treatment and animal setup habituation

At the end of the procedure, an iodine-based ointment (Braunodivon, 10%, B. Braun, Melsungen, Germany) was applied to the edges of the wound and a long-term analgesic (Meloxicam, 2 mg/kg, sc, Böhringer Ingelheim, Ingelheim, Germany) was administered and for 3 consecutive days. For at least 5 days post-surgery, the animal’s health status was assessed via a score sheet.

After at least 1 week of recovery, animals were gradually habituated to the experimental setup by first handling them and then simulating the experimental procedure. To allow for virus expression, neural recordings started after an incubation time of 2-4 weeks after injection.

### Craniotomy

On the day prior to the first day of recording, mice were fully anesthetized using the same procedures as described for the initial surgery, and a craniotomy (ca. 2×1 mm on the AP×BL axes) was performed over dLGN (ca. 2.5 mm posterior from bregma and 2.3 mm lateral from midline) and V1 and re-sealed with Kwik-Cast (WPI Germany, Berlin, Germany). As long as the animals did not show signs of discomfort, the long-term analgesic Metacam was administered only once at the end of surgery, to avoid any confounding effect on experimental results. Recordings were performed daily and continued for as long as the quality of the electrophysiological signals remained high.

### Extracellular multi-electrode array (MEA) recordings

After 2-4 weeks of incubation time, we performed *in vivo* extracellular multi-electrode array (MEA) recordings of dLGN neurons in awake, head-fixed mice (**Figure 1a**). Extracellular signals were recorded at 30 kHz (Blackrock microsystems, Blackrock Microsystems Europe GmbH, Hanover, Germany). For each recording session, the silicon plug sealing the craniotomy was removed. For dLGN recordings, a 32 channel linear silicon probe (Neuronexus A1×32Edge-5mm-20-177-A32) was lowered to a depth of ∼ 2500–3500 *µ*m below the brain surface. We judged recording sites to be located in dLGN based on the characteristic progression of RFs from upper to lower visual field along the electrode shank (*51*), the presence of responses strongly modulated at the temporal frequency of the drifting gratings (F1 response), and the preference of responses to high temporal frequencies (*56, 51*). For *post hoc* histological reconstruction of the recording site, the electrode was stained with DiI (Invitrogen, Carlsbad, USA) for some (typically the last) recording sessions.

### Locomotion

During the experiment, mice were free to run on an air-floating Styrofoam ball and the run speed was recorded via locomotion sensors (**Figure 1a**). Two optical computer mice interfaced with a microcontroller (Arduino Duemilanove) sampled ball movements at 90 Hz.

To compute animal run speed, we used the Euclidean norm of three perpendicular components of ball velocity (roll, pitch and yaw) and smoothed traces with a Gaussian kernel (*σ* = 0.2 s). To quantify the effect of running vs. sitting on various response properties, the run modulation index (*MI*_*Run*_) was defined based on the mean firing rates during running vs. sitting periods as

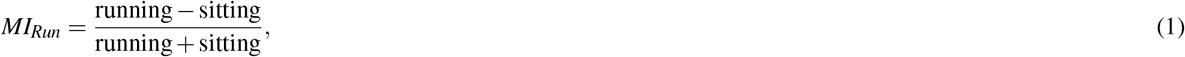

where running periods were defined as those for which speed exceeded 1 cm/s, and sit periods as those for which speed fell below 0.25 cm/s.

To test for a significant difference in mean FRs between the run vs. sit conditions matched for each neuron, we used the Wilcoxon signed-rank test.

### Eye tracking

To record eye position and pupil size, the animal’s eye that was viewing the stimulus was illuminated with infrared LED light and monitored using a zoom lens (Navitar Zoom 6000) coupled with a camera (Guppy AVT camera; frame rate 50 Hz, Allied Vision, Exton, USA).

Pupil position was extracted from the eye-tracking videos using a custom, semi-automated algorithm. Briefly, each video frame was equalized using an adaptive bi-histogram equalization procedure, and then smoothed using median and bilateral kernels. The center of the pupil was detected by taking the darkest point in a convolution of the kerneled image with a black square. Next, the peaks of the image gradient along lines extending radially from the center point were used to define the pupil contour. Lastly, an ellipse was fit to the contour, and the center and area of this ellipse was taken as the position and size of the pupil, respectively. A similar procedure was used to extract the position of the corneal reflection (CR) of the LED illumination. Eye-closure, grooming, or implausible ellipse fitting was automatically detected and the adjacent data points 0.15 s before and after were excluded. Linear interpolation and a subsequent Gaussian smoothing (*σ* = 0.06 s) was applied to fill the removed segments. Adjustable algorithm parameters, such as the threshold of the mean pixel-wise difference between each frame and a reference frame to detect blinks, were set manually for each experiment.

To quantify the effect of large vs. small pupil sizes on various response properties, the eye modulation index (*MI*_*Pupil*_) was defined based on the mean firing rates during periods of large vs. small pupils as

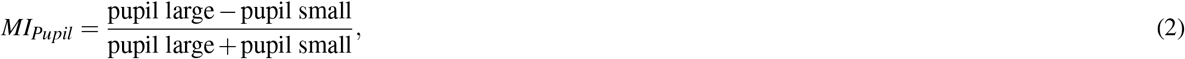

where periods of large pupils were defined as those for which pupil size was above the 50^th^ percentile of the median normalized pupil trace, and periods of small pupils as those for which pupil size fell below the 25^th^ percentile.

### Optogenetic feedback suppression

To photosuppress V1 Ntsr1+ L6 CT pyramidal cells, an optic fiber (480 *µ*m core diameter, MFP_480/500/1000-0.63_m_SMA, Doric Lenses, Quebec, Canada) was coupled to a light-emitting diode (blue LED, center wavelength 465 nm, LEDC2_465/635_SMA, Doric Lenses, Quebec, Canada) and positioned with a micromanipulator less than 1 mm above the exposed surface of V1. A black metal foil surrounding the tip of the head bar holder prevented the photostimulation light from reaching the animal’s eyes. To ensure that the photostimulation was effective, the first recording session for each mouse was carried out in V1. Only if the exposure to light reliably induced suppression of V1 activity was the animal used for subsequent dLGN recordings. LED light intensity was adjusted on a daily basis to evoke reliable effects and account for variations in exact virus titer, volume, incubation time, virus expression levels, and fiber position (0.85-9.5 mW at the fiber tip). Since the tip of the fiber never directly touched the surface of the brain, and since the clarity of the surface of the brain varied (generally decreasing every day following the craniotomy), the light intensity delivered even to superficial layers of V1 was inevitably lower. For the movie stimulus, optogenetic pulses of 1 s duration were sent randomly each second with a 50 % chance.

To quantify the effect of CT feedback suppression on various response properties, we defined the optogenetic modulation index (*MI*_*CT FB supp*._) based on the mean FRs during CT feedback suppression (‘opto’) versus the control condition as

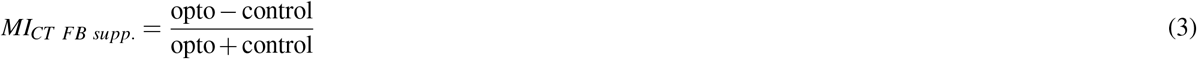

### Joint modulation by additional inputs

In order to quantify the joint effect of all the modulatory inputs, we calculated *MI*_*Joint*_ as

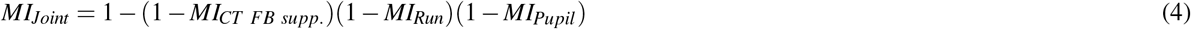

### Visual stimulation

During the experiment, the mice were passively viewing visual stimuli on an LCD monitor screen in their right visual field. The visual stimuli were presented on a gamma-calibrated liquid crystal display (LCD) monitor (Samsung SyncMaster 2233RZ, 47×29 cm, 1680×1050 resolution at 60 Hz, mean luminance 50 cd/m^2^) positioned at a distance of 25 cm from the animal’s right eye (spanning ∼ 108×66° visual angle by small angle approximation) using custom written software (EXPO, https://sites.google.com/a/nyu.edu/expo/home). The display was gamma-corrected for the presentation of artificial stimuli, but not for movies (see below).

#### Movie stimulus

For movie stimulus generation, we adopted a set of randomly picked clips from various movies. Briefly, source movie clips were converted to grey scale, temporally downsampled to 30 frames per s, spatially resampled and cropped to 424×264 pixels, to be presented on our 47×29 cm monitor screen at 25 cm distance at 106×66° (4 pixels/°) visual angle (by small angle approximation, which preserves the desired pixel resolution at the screen center better than the arctangent). Movie frames were not histogram-equalized and presented at 60 Hz (repeating each frame twice) without monitor gamma correction, since cameras are already gamma corrected for consumer displays (*105*). To generate the movie sequence, we used a random set of 296 unique movie clips (5 s each) and split 188/296 clips into 8 parts of 36 unique clips (5 s×36 = 180 s per part). They were interleaved with set of 8 clips (5 s×8 = 40 s) which was repeated 9 times. The repeated clips served to give an estimate of response reliability to the same clips. The movie sequence was flanked by a period of blank grey screen presentation (1 min) at the beginning and at the end, to record spontaneous activity. This resulted in a total stimulus duration of ∼ 32 mins. To rule out sequence effects, we randomized the clip order for different stimulus presentations. To investigate the effects of L6 CT FB suppression, we simultaneously presented a random optogenetic pulse train of 1 s pulses, occurring each second with a probability of 50 %, throughout the entire stimulus duration, including blank grey screen periods.

#### Sparse noise stimulus

To measure RFs in a more standard manner, we also presented an (artificial) sparse noise stimulus. The stimulus consisted of a rapid sequence of non-overlapping white and black squares appearing in succession within a 12×12 square grid presented on a grey background of mean luminance (50 cd/m^2^). The square grid spanned 60° per side, while individual squares spanned 5° per side. Each square flashed 20 times for 200 ms at random order. The stimulus triggered average (STA) for the sparse noise stimulus was computed using the onset of each square and then computing the normalised mean spike rate triggered by each position.

### Histology

To verify virus expression and recording sites, we performed post-mortem histological analyses. After the final recording session, mice were first administered an analgesic (Metamizole, 200 mg/kg, sc, MSD Animal Health, Brussels, Belgium) and following a 30 min latency period were transcardially perfused under deep anesthesia using a cocktail of Medetomidin (Domitor, 0.5 mg/kg, Vetoquinol, Ismaning, Germany), Midazolam (Climasol, 5 mg/kg, Ratiopharm, Ulm, Germany) and Fentanyl (Fentadon, 0.05 mg/kg, Dechra Veterinary Products Deutschland, Aulendorf, Germany) (ip). Perfusion was first done with Ringer’s lactate solution followed by 4% paraformaldehyde (PFA) in 0.2 M sodium phosphate buffer (PBS). Brains were removed, postfixed in PFA for 24 h, and then rinsed with and stored in PBS at 4° C. Slices (50 *µ*m) were cut using a vibrotome (Leica VT1200 S, Leica, Wetzlar, Germany), stained with DAPI-solution (DAPI, Thermo Fisher Scientific, Waltham, Massachusetts, USA), mounted on glass slides with Vectashield mounting medium (Vectashield H-1000, Vector Laboratories, Burlingame, USA), and coverslipped. A scanning fluorescent microscope (BX61, Olympus, Tokyo, Japan) was used to inspect slices for the presence of red fluorescent protein (RFP/FusionRed) marking stGtACR2-channels, and DiI, marking electrode tracks. Recorded images were processed off-line using FIJI (*106, 107*).

### Control experiments with Ntsr-negative mouse

Additionally, same experimental procedures were followed for control (Ntsr-negative) mouse, in order to assess the efficiency of optogenetic manipulations of CT feedback. Prior to experimentation, the genotype of the mice was confirmed via polymerase chain reaction (PCR) analysis.

### Spike sorting and unit extraction

Spike sorting was performed to obtain single unit activity from extracellular recordings. Electrophysiological signal recordings were kerneled using a 4^th^-order Butterworth high-pass non-causal kernel with a low frequency cutoff of 300 Hz. We then used the open source, MATLAB-based (The Mathworks, Natick, Massachusetts, USA), automated spike sorting toolbox Kilosort and Kilosort2 (*108*). Resulting clusters were manually refined using Spyke (*109*), a Python application that allows for the selection of channels and time ranges around clustered spikes for realignment, as well as representation in 3D space using dimension reduction (multichannel PCA, ICA, and/or spike time). In 3D, clusters were then further split via a gradient-ascent based clustering algorithm (GAC) (*110*). Exhaustive pairwise comparisons of similar clusters allowed the merger of potentially over-clustered units. For subsequent analyses, we inspected autocorrelograms and mean voltage traces, and only considered units that displayed a clear refractory period and a distinct spike waveshape.

### Data analysis

Data analysis was performed using custom-written code that applies general tools such as Numpy, sklearn, matplotlib, seaborn, pandas and carried out in a MySQL-based database using the DataJoint (*111*). We also used a customized version of RFEst (https://github.com/berenslab/RFEst) to allow for multiple model inputs.

### Firing rate calculations

To obtain units firing rates in spikes per second (Hz), each unit’s spike density function (SDF) was calculated by binning spikes into a firing rate histogram (bin width = 20 ms) and convolving this with a Gaussian of width 2*σ* = 10 ms. Mean firing rates (FRs) over a given condition were calculated as the mean of the time-varying firing rates for the defined periods.

### Identification of putative neuronal types

In order to identify putative excitatory and inhibitory neurons in V1, we analysed the extracellular spike waveform. For each neuron, the mean waveform of the maximally responsive electrode channel was obtained, and the time between trough and peak (trough-to-peak time) and the full-width at half-height of the peak (peak width) were calculated. Using the waveforms of all V1 neurons, a k-means algorithm was used to cluster the data into 2 populations.

### Response reliability

To compute how reliable a neurons responded to the visual stimulus we used the set of 8 clips that where repeated 9 times throughout the experiment. We computed reliability by correlating each repetition with the mean of all other repetitions and averaging that over all splits.

### Optogenetic significance

To compute the reliability of the optogenetic manipulation of L6 CT neurons, we developed a trial-based permutation test. In response to drifting gratings, we calculated the average firing rate during each trial and separated the trial with optogenetic CT feedback suppression (*trials*_*CT FB supp*._; n=90 − −130) from those without (*trials*_*control*_; n=90 − −130). We calculated the observed statistics *e f f ect*_*CT FB supp*._ as the difference in the means between the *trials*_*CT FB supp*._ and the *trials*_*control*_. We assessed significance by permuting trial labels 1000 times and considered the effect significant if it fell outside of the distribution of permuted *e f f ect*_*CT FB supp*._s. Finally, we compared the percentage of neurons that passes the significance test between the included recording and a recording from a control mouse (Ntsr-negative).

### Exclusion criteria

Neurons with mean evoked firing rates < 0.1 Hz were excluded from all further analysis. Further analysis-specific selection criteria are stated in the appropriate subsections.

### Predictor correlations

To test for the correlations between the predictors *stimulus, opto, run, and eye*, we temporally aligned these traces, including only time points for which we had data points in all traces (e.g. removing periods of eye blinks). We then explored their cross-correlations in order to detect potential delays in their effects on each other. Such is the case, for instance in the pupil light reflex, where increases in stimulus intensity are followed by a delayed decrease in pupil size. We then used the delay time to shift the traces appropriately before computing their correlation value (Pearson’s *r*; **Figure S3c**).

In order to test for statistical significance of the obtained correlation values, we first needed to account for the fact that our time-series inherently contain autocorrelations which would lead to an overestimation of correlations between them (except for the random opto pulses) (*53*). We therefore used a permutation test in which we randomly permuted the stimulus traces for *k* = 1000 iterations, and then computed the p-value of the observed correlation value as its percentile within the null distribution of p-values for the permuted traces (*53*).

### Spline-based generalized linear model

To estimate the spatio-temporal RFs (STRFs), we used the RFEst Python toolbox for spline-based spatio-temporal RF estimation (*38*). Here, the spline-based GLM reduces the number of parameters compared to traditional approaches, which need to estimate every pixel in the RF independently, while spline-based GLM only estimate the weight of the bases. RFEst is also less data demanding and reduces the computation time significantly (*38*). The number of parameters are given by the number of basis functions, also referred to the degrees of freedom. By using natural cubic splines as the basis (e.g., **Figure S6a–d**), the estimates are automatically smooth, which is a desirable property for single STRFs. To impose sparsity on the weights (also a desirable property of SRFs) we added L1 regularization, which pushed the weights for less relevant bases to zero. To compute the spline-based STRFs, *w*_*SPL*_, the coefficients, *b*_*SPL*_, were obtained as

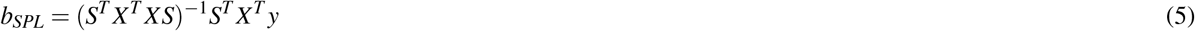

with *X* as the stimulus design matrix, *y* as the neural response vector, and *S* as the spline matrix. The spatio-temporal RF was computed as

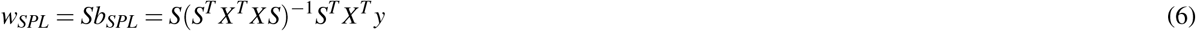

To generate the natural cubic spline matrices (*S*) the package Patsy (0.5.1) is integrated into RFEst.

To approximate *w*_*SPL*_ we used a generalized linear model (GLM) that predicts the instantaneous firing rate for one neuron using the movie as a predictor. We extended this “Stimulus only” model by integrating also running speed and pupil size as behavioral predictors of neuronal firing rate. To estimate the effect of cortico-thalamic (CT) feedback, an additional bimodal input was used comprising the optogenetic light stimulation that could be either on (*o*_*t*_ = 1) or off (*o*_*t*_ = 0). All inputs where parameterized with a set of spline basis and multiplied with an extra weight vector (also referred to as kernel):

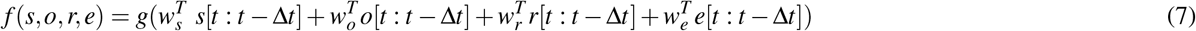

with *s, o, r*, and *e* denoting the additional model inputs of stimulus, optogenetics, running and eye, respectively, and [*t* : *t* − Δ*t*]defining the temporal integration window (250 ms for stimulus and 800 ms for the predictors for modulatory inputs).

We estimated the kernels by gradient descent with respect to the cost function with L1 regularization. As standard procedure for time series data, we used 150 out of the 188 unique movie clips (80 %) for cross-validation to select optimal hyperparameters and reserved the remaining 38 clips (20 %) as a held-out test set. To select optimal hyperparameters, we used five-fold cross-validation grid search on the training data (120 training clips and 30 validation clips in each fold). Hyperparameters included the number of spline basis (between 10 and 19) in temporal dimension (for stimulus, pupil size, locomotion, and feedback input) and spatial dimension (only stimulus), as well as the strength of the L1 regularization (with weights varying from 5 to 15). The STRFs were initialized randomly and optimized using gradient decent for 2000 iterations. We stopped the fitting early when the training cost changed less than 10^−5^ for 10 iterations. Finally, we selected the hyperparameters based on the mean performance on the validation set across folds. After optimal hyperparameter selection, models were retrained on the full training data (150 movie clips) and the final performance of the model was reported as the correlation coefficient between predicted and observed neural responses on the held out test data (38 movie clips).

We further fitted spline GLMs with responses to a sparse noise stimulus. We conducted a separate hyperparameter search for this stimulus, using a similar procedure as for the movie stimulus, involving cross-validation. For each neuron, we determined a unique set of optimal hyperparameters and kernels for the sparse noise stimulus and the movie stimulus.

### Session based permutation test

Following Harris (2020) (*53*), we performed a permutation test to evaluate if data from a different experimental session would lead to the same or worse model performance, which indicated the significance of the input-output correlations captured by the model. To achieve this, we provided the model with input data from the validation set of an unrelated session (for the model inputs ‘stimulus’, ‘running’, ‘pupil size’) or with synthetic inputs generated with the same statistics as the original (for the ‘CT feedback suppression’ input), one at a time. We repeated the process using all different data from all recording sessions. Subsequently, we compared for each input the actual model performance (Pearson’s *r*) on the validation set across its folds (n=5) against a distribution of model performances with that specific input permuted from the different recording sessions across their folds (n=5 folds × 9 recording sessions). Inputs were considered significant if the actual performance differed from the permuted performance with *p* ≤ 0.05 using the non-paired Mann-Whitney-U test.

### Spatio-temporal RF component extraction

To separate spatial and temporal components of the 3D STRFs, we performed singular value decomposition (SVD) on the norm of the stimulus weight vector *w*. The temporal RF is extracted as the first left-singular vector of *U*, i.e. temporal dimension with the highest variance, and the spatial RF as the first right-singular vector of *V*, reshaped into the height- and width-dimensions of the input vector *w*. The extremes of the reshaped spatial RF vector are then used to quantify RF position and RF area. The extracted temporal RF components were normalized and multiplied with the RF center value before computing the slope (−150 ms to peak).

### Spatial RF contour estimation

Model spatial RFs were estimated by extracting the 2D spatial RF component from the model weights (see subsection ‘Spatio-temporal RF component extraction’), and then drawing a contour line around the largest absolute peak (assumed to be the center of the spatial RF). The contour threshold gets gradually lowered until any further decrease would result in a second contour around the second largest extremum (background irregularities considered as noise). To avoid overly large RFs in very clean spatial components (without any major second extremum), the contour threshold had to be 2 standard deviations above or below the mean. To improve estimate accuracy, the spatial RF component was upsampled 16-fold via cubic spline interpolation.

### Spatial RF area

The spatial RF areas were estimated by using the spatial RF component and contour (see subsection ‘Spatial RF contour estimation’) and calculating the number of pixels of the spatial RF contour mask in relation to the total number of pixels in the image frame, which was then scaled by the stimulus extent to obtain the value in squared degrees of visual angle. To improve estimate accuracy, the spatial RF component was upsampled 16-fold via cubic spline interpolation.

### Spatial RF center-surround

stRFs were collapsed across the azimuth axis, and accordingly defined as a function of elevation and time. Then, we identified the time point with peak activity. Subsequently, centre regions were defined according to the spatial width of the peak activity, and surround regions were defined as a ring encircling the centre and extending up to trice the diameter or 9^*°*^. We then summed pixel intensities within the centre region and the surround regions separately and calculated surround-to-centre ratio as:

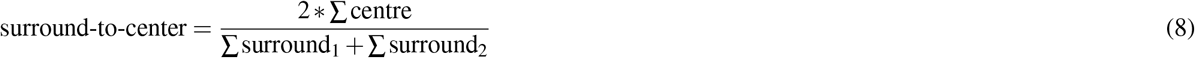

### Functional cell types clustering

To categorize neurons into functional subtypes, we employed dimensionality reduction (PCA) on their PSTH responses to a full-field light intensity step stimulus. Subsequently, we conducted clustering using a Gaussian mixture model on the principal components, resulting in the identification of four primary groups (Sustained ON, Sustained OFF, Transient, and Mixed). Nevertheless, we observed substantial response diversity even within these main groups. To address this, we further subdivided neurons within each group through empirical selection of cluster numbers, aiming to optimize silhouette scores and reduce the standard error of the PSTH mean response within the subclasses. This two-step clustering approach allowed capturing finer distinctions of response patterns (**Figure S5a, b**).

### Rate of change (RoC) to CT feedback suppression

To quantify the temporal dynamics of the effect of suppressing CT feedback with optogenetic pulses, we calculated rate of change (RoC) in the neurons’ responses based on both data and model prediction. For each case, we first identified the time point *t*_*min*_ where the slope of neurons response flipped its sign (e.g. after light onset, firing rates decreased, negative slope, and then it either stays constant or fluctuates around its new value and the slope becomes zero or flips its sign to positive). Then, we calculated the normalised change in responses *MI*_*CT FB supp*._; (see subsection ‘Optogenetic feedback suppression’). Finally, *RoC* was defined as:

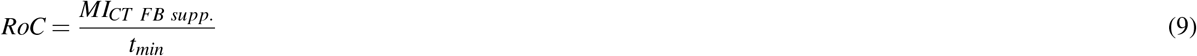

### Decoding analysis

We used a two-alternative forced choice (2AFC) decoder (*30*) to analyze if stimulus discriminability is easier with or without feedback. Given two spike trains *{s*_*A*_, *s*_*B*_*}* in response to two movie clips *{A, B}* we computed the model log-likelihood for the correct ((*s*_*A*_, *A*), (*s*_*B*_, *B*)) and the incorrect pairing ((*s*_*A*_, *B*), (*s*_*B*_, *A*)). We used the ‘Full model’ to predict spike trains in the two feedback conditions, (1) with the feedback component intact and (2) with the feedback component suppressed. In both conditions, we used 100 randomly selected 50 ms movie clips from the test set and their corresponding responses. Using all possible pairs, we computed the model log-likelihood for the correct and the incorrect pairing in both feedback conditions. The log probability for the correct pairing of response *s*_*A*_ and stimulus *A* is defined as follows:

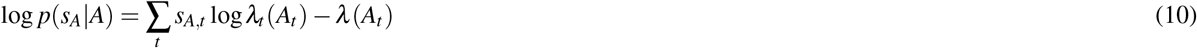

where *λ* is the instantaneous firing rate at time *t* predicted by the model. Analogously, we computed log *p*(*s*_*B*_|*B*), log *p*(*s*_*A*_|*B*), and log *p*(*s*_*B*_|*A*). A correct choice was made if log *p*(*s*_*A*_|*A*) > log *p*(*s*_*A*_|*B*) or log *p*(*s*_*B*_|*B*) > log *p*(*s*_*B*_|*A*), respectively. We used the percentage correct over all possible pairs to quantify decoding performance. Finally, we computed the ratio of percentage correct for the feedback suppressed condition and the control condition. A ratio above one indicates better decoding performance in the feedback suppressed condition and a ratio below one in the control condition.

## Supplementary Materials

**Figure S1.**
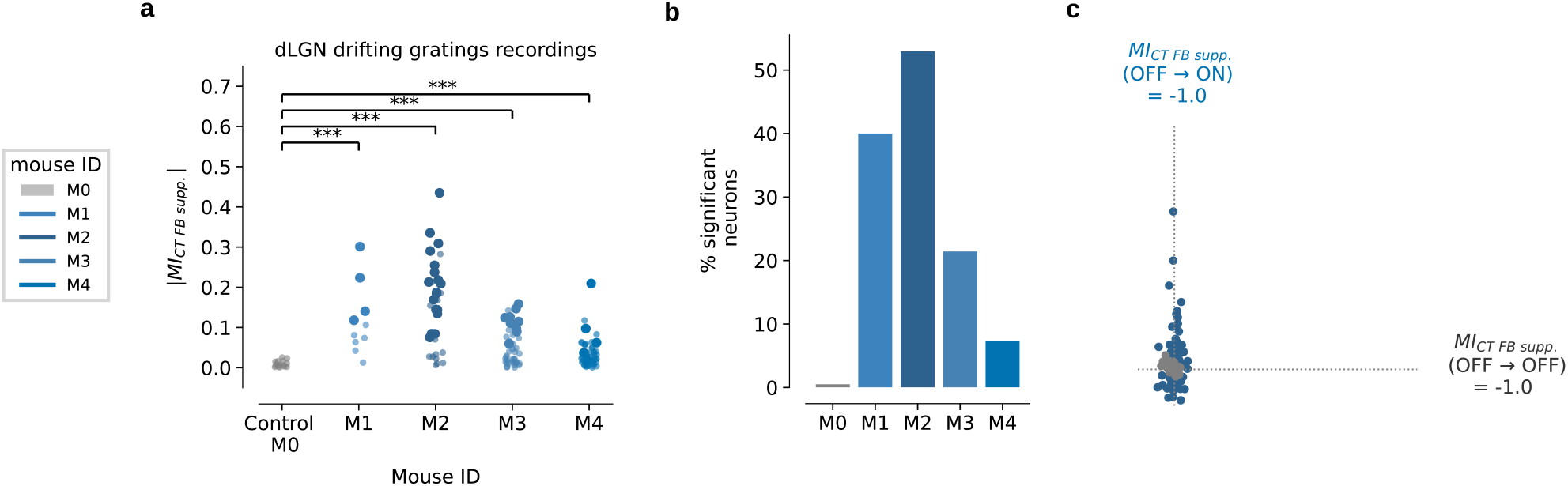
Efficiency of optogenetic manipulation of CT feedback. (**a**) Optogenetic modulation index (OMI) values for dLGN neurons (n = 122), separately by mouse. The four included mice showed significantly higher modulation indices then the control mouse M0 (*p* < 0.01, Mann-Whitney-U test). *Opaque dots*: Neurons that are significantly modulated by CT feedback suppression (*p* < 0.05, trial-based permutation test). *Transparent dots*: Neurons that did not pass the significance test (*p* > 0.05). (**b**) Percentage of neurons that were significantly modulated by CT feedback suppression per mouse (*p* < 0.05, trial-based permutation test). (**c**) Scatter of OMI values for the four included mice (*blue*) and the control mouse (*Gray*) for the OFF-ON transitions (i.e., triggered to the onset of CT feedback suppressing light pulses) vs. OFF-OFF transitions (i.e., triggered to times without onset of CT feedback suppressing light pulses).

**Figure S2.**
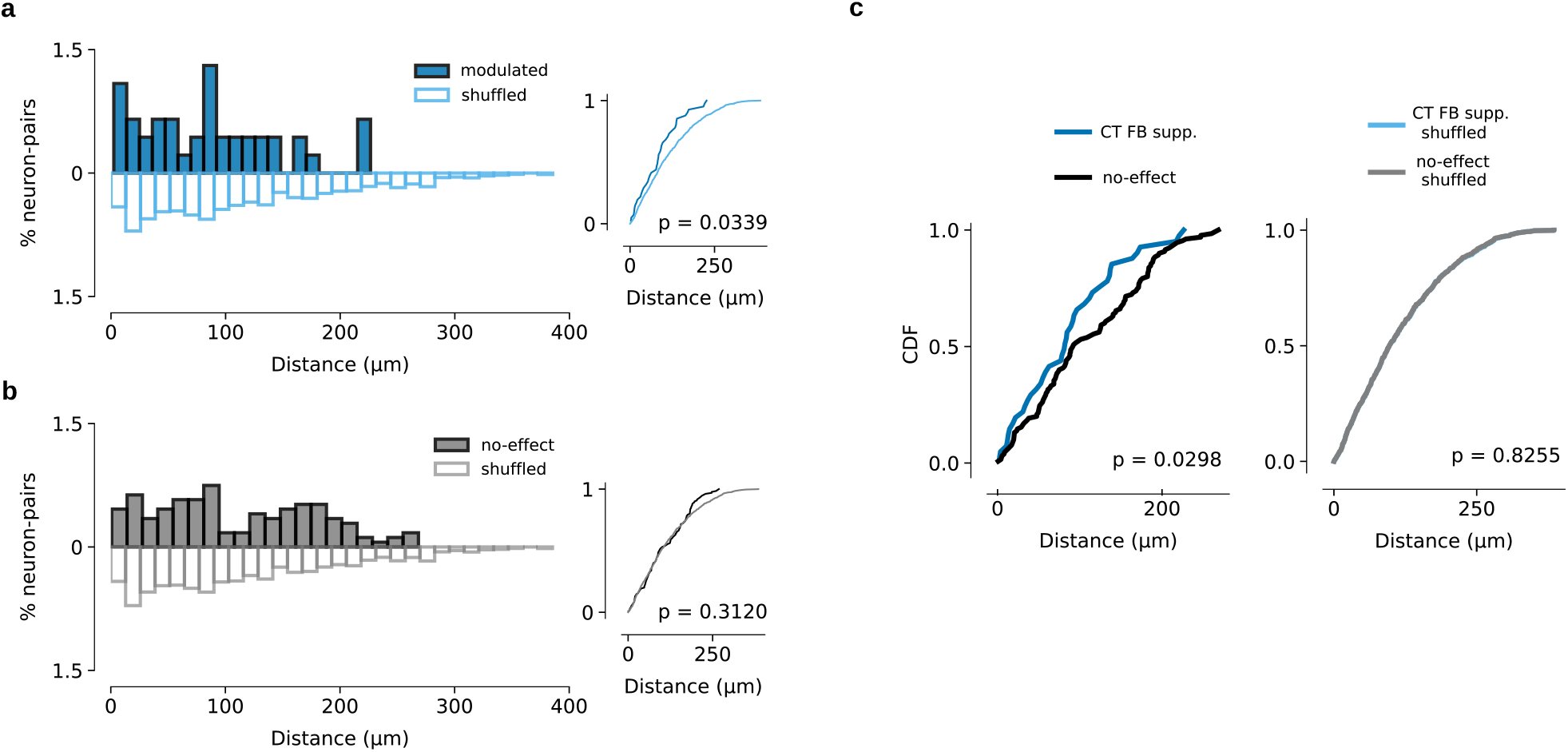
CT feedback modulated neurons are closer to each other in the mouse dLGN. We identified all simultaneously recorded neuron pairs (*n* = 171), of which at least one neuron was positively or negatively modulated by CT feedback suppression (|*MI*_*CT FB supp*._| ≥ 0.1), and calculated the distance in micrometer between each pair on the electrode. (**a**) *Left, solid bars*: Percentage of neuron pairs (*n* = 41) at each distance from each other on the electrode, when both neurons were modulated by CT feedback suppression. *Left, hollow bars*: Percentage of the same neuron-pairs with shuffled distances. *Right*: cumulative distributions of true and shuffled data(*p* = 0.0339, Kolmogorov-Smirnov test). (**b**) Same as (a), when only one neuron was modulated by CT feedback suppression (*n* = 130). *p* = 0.312, Kolmogorov-Smirnov test. (**c**) *Left*: Cumulative distributions of distances between neuron pairs(*p* = 0.0298, Kolmogorov-Smirnov test). *Right*: cumulative distributions of shuffled data (*p* = 0.8255, Kolmogorov-Smirnov test), re-plotted from (a) and (b).

**Figure S3.**
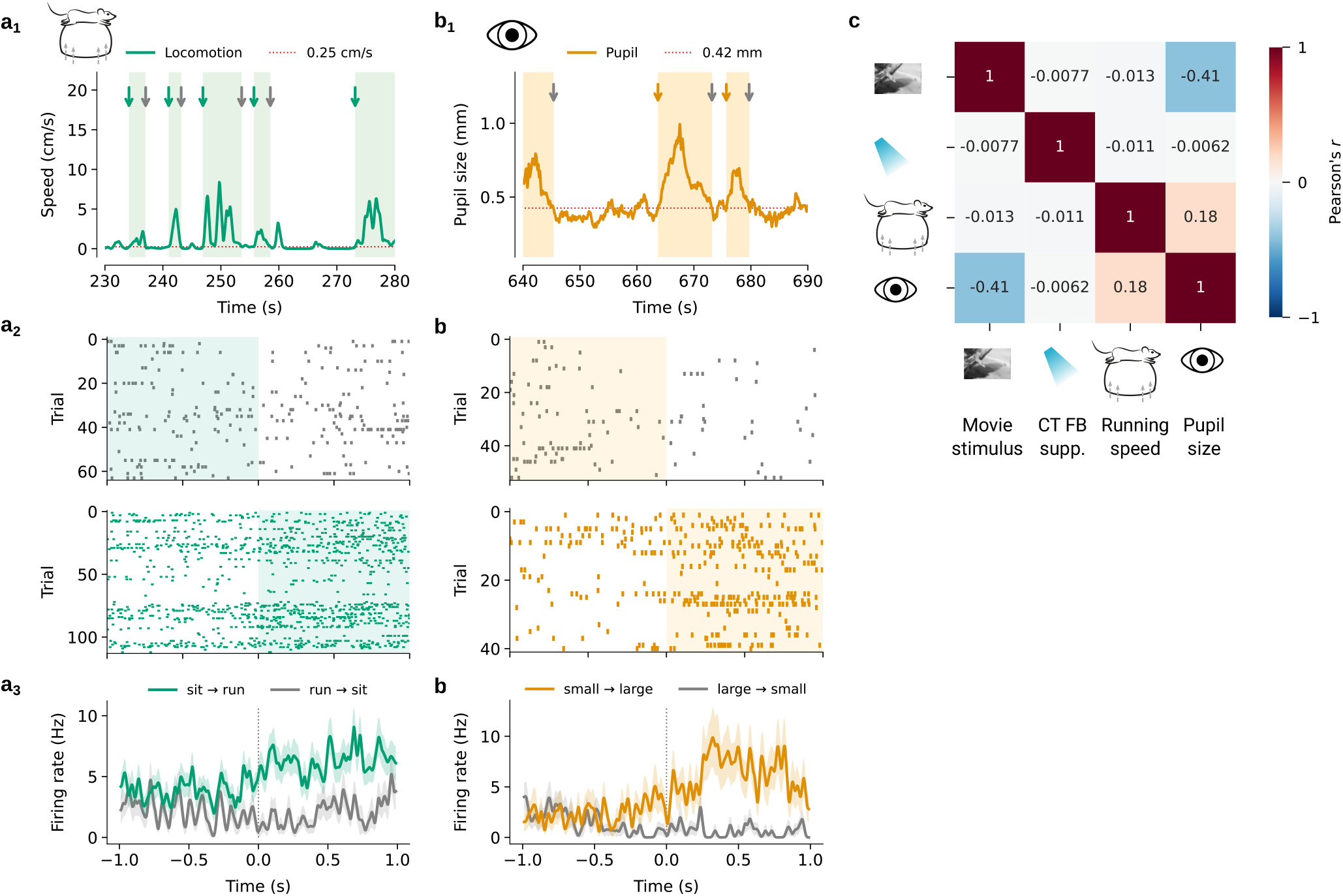
CT feedback and behaviour modulates dLGN responses to movie clips. (**a**) Modulation of firing rate by running speed for one dLGN example neuron: (a_1_) We considered periods as “running” if the animal’s speed was > 1 cm/s, and “sitting” if the speed was < 0.25 cm/s (*red*). Transitions between periods are marked by green and grey arrows, respectively. (a_2_) Spike raster plot triggered on transitions from running to sitting (*top*), and sitting to running (*bottom*). (a_3_) Corresponding PSTHs. *Green*: transitions from sitting to running; *grey*: transitions from running to sitting. Only periods with durations > 2 s were considered. (**b**) Modulation of firing rate by pupil size for one dLGN example neuron: (b_1_) We considered periods as “large pupil size” when pupil size exceeded the median pupil size during the experiment (*red*), and “small pupil size” if it was smaller than the 25th percentile. Transitions between periods are marked by orange and grey arrows, respectively. (b_2_) Same as in a_2_ but for transitions from large to small pupil (*top*) and from small to large pupil (*bottom*). (b_3_) Same as in a_3_ but for pupil size transitions. (**c**) Correlation matrix of average stimulus intensity, optogenetic light pulses, running speed and pupil size traces. Values denote average correlation values across experiments (*n* = 10).

**Figure S4.**
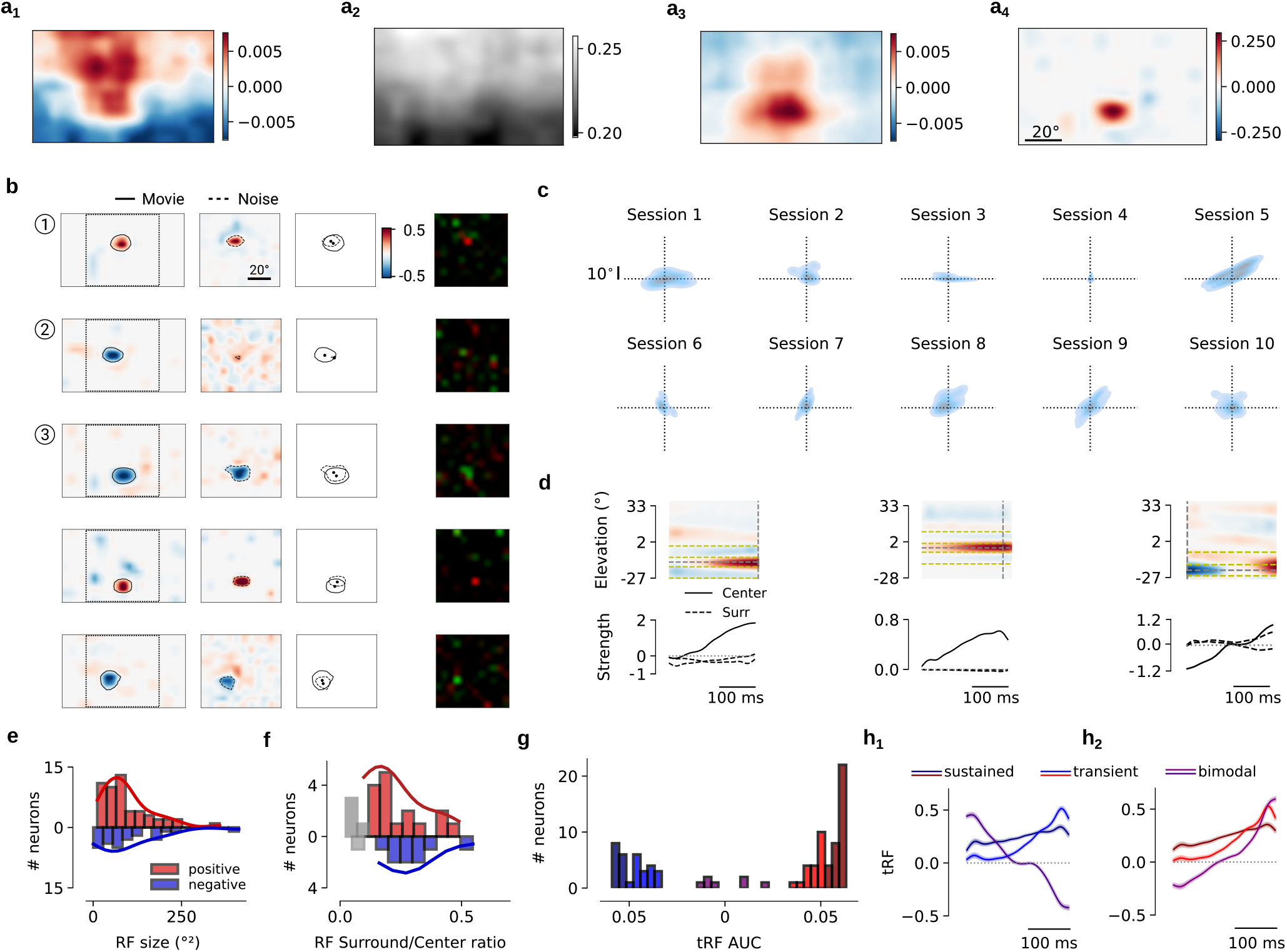
Spatial RF estimation by the spline-GLM model. (**a**) Comparison of spatial RF (sRF) estimates during the movie stimulus for one example dLGN neuron. (a1) Simple spike triggered average (STA). (a2) Pixel intensity averaged across all movie frames, showing a gradient from top to bottom typical for naturalistic scenes. (a3) STA corrected for average intensity gradient. This yields a RF with similar centre and polarity compared to the one learned by the spline-GLM (a4), but with larger and more diffuse area. (a4) Learned sRF by the spline-GLM model. (**b**) To validate the sRF on a neuron-by-neuron basis, we compared the spline-GLM sRF learned from movies (*first column*) with spline-GLM sRF learned from sparse noise experiments (*second column*; overlay of the two sRF centres and contours in the *third column*; see Methods) and with the stimulus-triggered average response map for the sparse noise experiments computed independently from the model (*last column*). The first three example neurons are the same as in **Figure 2c**. (**c**) Centre shift between movie and sparse noise sRFs for all 10 experimental sessions (neurons with significant stimulus kernels, *n* = 85). Position around the origin indicates close to zero shift. (**d**) Spatio-temporal RF (stRF) for three example neurons. *Top*: stRFs collapsed across the azimuth axis, and plotted as a function of elevation and time. *Vertical dashed line*: time point with peak activity. *Horizontal dashed lines*: centre regions defined according to the spatial width of the peak activity, and surround regions defined as a ring encircling the centre and extending up to trice the diameter or 9^*°*^.*Bottom*: stRF centre (*solid*) and surround (*dashed*) activity. *Left*: antagonistic surround neuron (surround-to-centre ratio = 0.32). *Middle*: non-antagonistic surround neuron (surround-to-centre ratio = 0.02). *right*: bi-phasic response neuron. (**e**) Area of sRF centres, separately for neurons with positive (*red*) and negative (*blue*) RF centre and significant stimulus kernels. Of these, ∼ 61%(52/85) had a positive centre, ∼ 29%(25/82) had a negative centre, and ∼ 9%(8/85) had a bimodal response where the detected RF changed its polarity across time.Median RF radius: 5 deg; median centre area: 81 deg^2^ for positive RF centers and 77 deg^2^ for negative RF centers. (**f**) Surround-to-centre ratios for the population of dLGN neurons with significant stimulus kernels, and antagonistic activity in the surround region (∼ 41%(35/85) neurons). We defined a threshold at 0.1 surround-to-centre ratio (*gray*). Of the remaining neurons with pronounced antagonistic surround, 16 neurons had positive RF centre with a negative surround (*red*), 9 neurons had negative RF centre with a positive surround (*blue*), and 4 neurons showed a biphasic response together with an antagonistic surround. (**g**) Quantification of tRF properties: histogram of area under the tRF, separately for neurons with positive (*red*) and negative (*blue*) RF centre. Neurons with small tRF AUC values ≦ 0.035 represent the bimodal neurons (*n* = 8, *purple*). Neurons with the largest tRF AUC values ≥ 0.052 were defined as neurons with sustained responses, while the remaining neurons were defined as neurons with transient responses. (**h**) Mean tRF split into sustained, transient, and biphasic.

**Figure S5.**
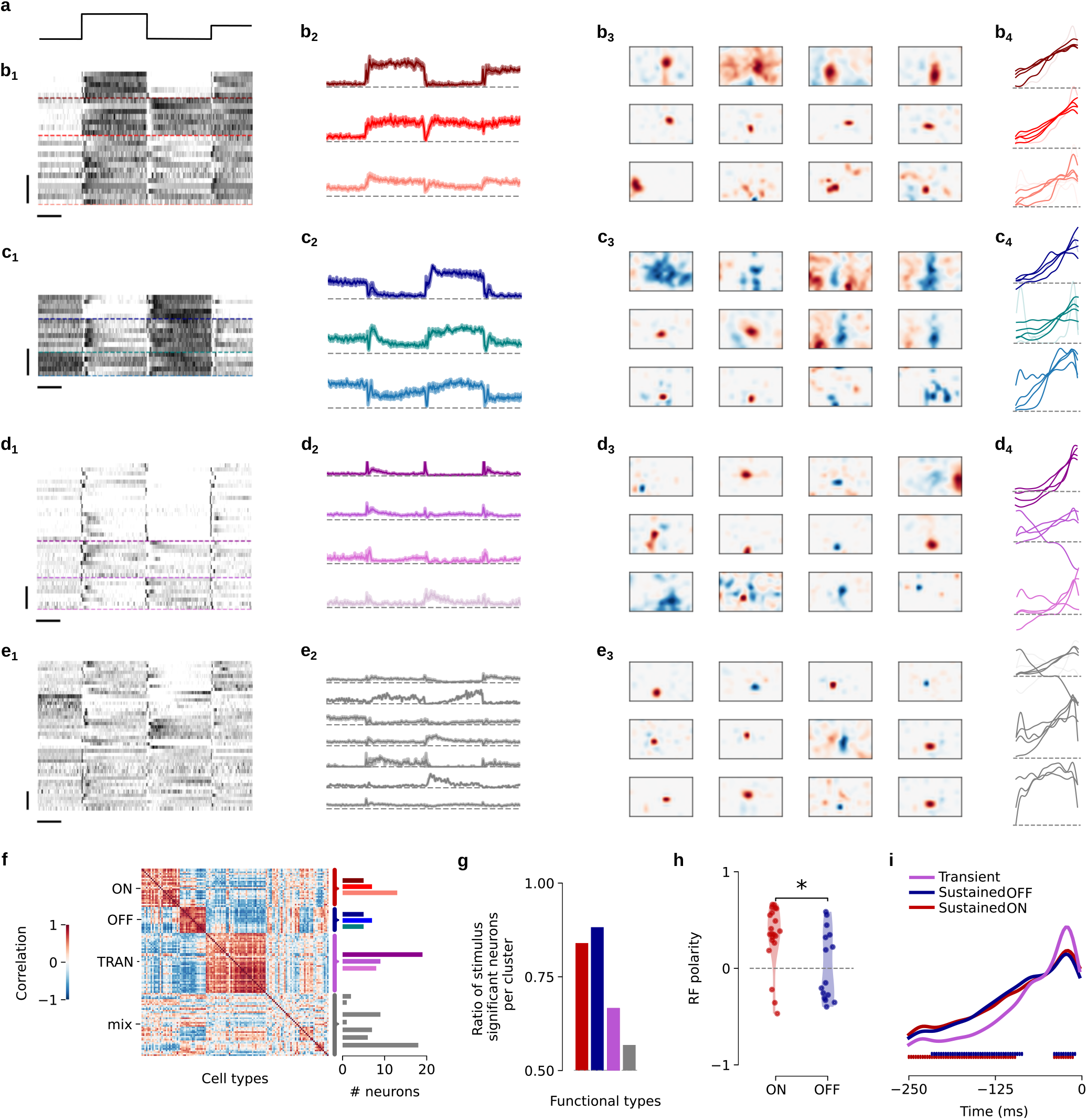
Spline-based GLM captures spatio-temporal RFs of different functional cell types in dLGN. (**a**) *Top, left*: Light intensity steps of the full field stimulus used for functional cell type classification. (**b**) Responses of all neurons assigned to the Sustained ON group (b_1_, *n* = 25), along with their average sub-cluster responses (b_2_). Spatial (b_3_) and temporal RF kernels (b_4_) of example neurons with significant movie stimulus kernel learned by the spline-GLM independently of the clustering. (**c**) Same as (b), for the Sustained OFF group (*n* = 17). (**d**) Same as (b), for the Transient group (*n* = 36). (**e**) Same as (b), for the remaining neurons (mixed cluster, *n* = 44). (**f**) *Left*: Correlation matrix of PCA features used for clustering. Neurons are sorted by groups. *Right*: Neuron counts per group. (**g**) Fraction of neurons with significant stimulus kernel in the spline-GLM model per functional type based on clustering. (**h**) Spatial RF polarity (sign of the spline-GLM fitted RF centre) for the ON and the OFF group based on clustering (*p* = 0.018, Mann-Whitney-U test). (**h**) Average temporal RF of movie kernels learned from spline-GLM for the OFF, ON, and Transient groups. *Horizontal bars*: Significant differences between OFF and Transient groups (*blue*) and between ON and Transient groups (*red*).

**Figure S6.**
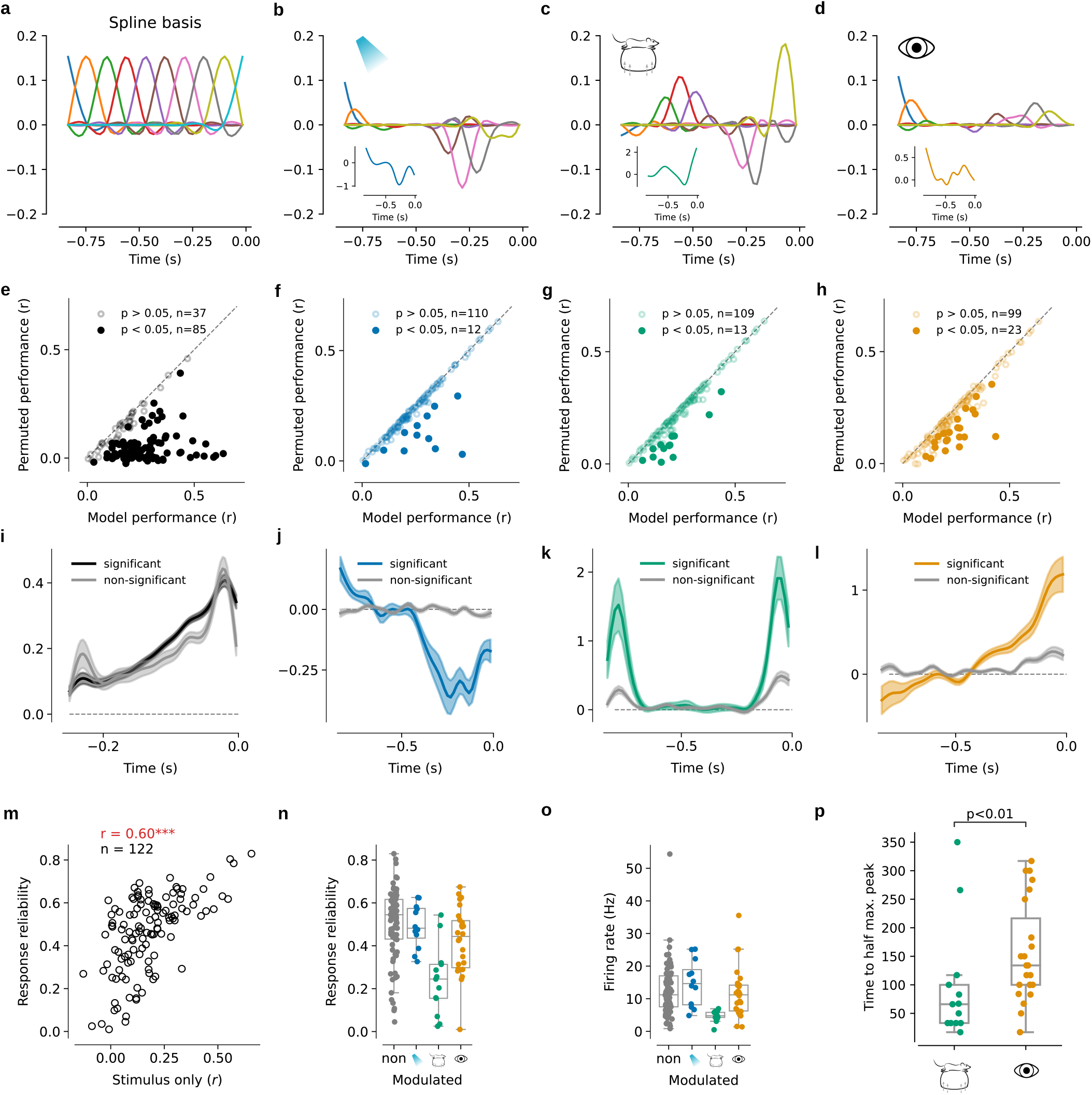
Spline-GLM model kernels for modulatory inputs. (**a**) Illustration of the natural cubic spline basis functions (n=10). (**b**) Weighted bases of the natural cubic splines after fitting the CT feedback kernel to the data for one example neuron. CT feedback kernel is shown in the inset. (**c**) Same as (b), for run speed kernel. (**d**) Same as (b), for pupil size kernel. (**e**) Model performance for permuted visual stimulus input (mean over all permutations) vs. actual performance. *Filled circles*: significant neurons. (**f**) Same as (a), for CT feedback suppression. (**g**) Same as (a), for running. (**h**) Same as (a), for pupil size. (**i**) Average temporal RF kernels, separately for neurons with significant (*black, n* = 85) versus non-significant (*grey, n* = 37) spatio-temporal kernels. *Shaded areas*: SEM. (**j**) Same as (e), for neurons with significant (*blue, n* = 12) versus non-significant (*grey, n* = 110) CT feedback kernels. (**k**) Same as (e), for neurons with significant (*green, n* = 13) versus non-significant (*grey, n* = 109) run kernels. (**l**) Same as (e), for neurons with significant (*orange, n* = 23) versus non-significant (*grey, n* = 99) pupil kernels. (**m**) Model performance versus response reliability across all neurons. Response reliability was computed as the mean correlation of dLGN responses to each 40 s repeated movie clip with all other 8 repeats. (**n**) Response reliability of dLGN neurons split into neurons with significant kernels to CT feedback suppression, running, pupil size, or non-modulated.(**o**) Same as (j) but for firing rate of dLGN neurons. (**p**) Kernel time to half maximum peak for run and pupil size kernels.

**Figure S7.**
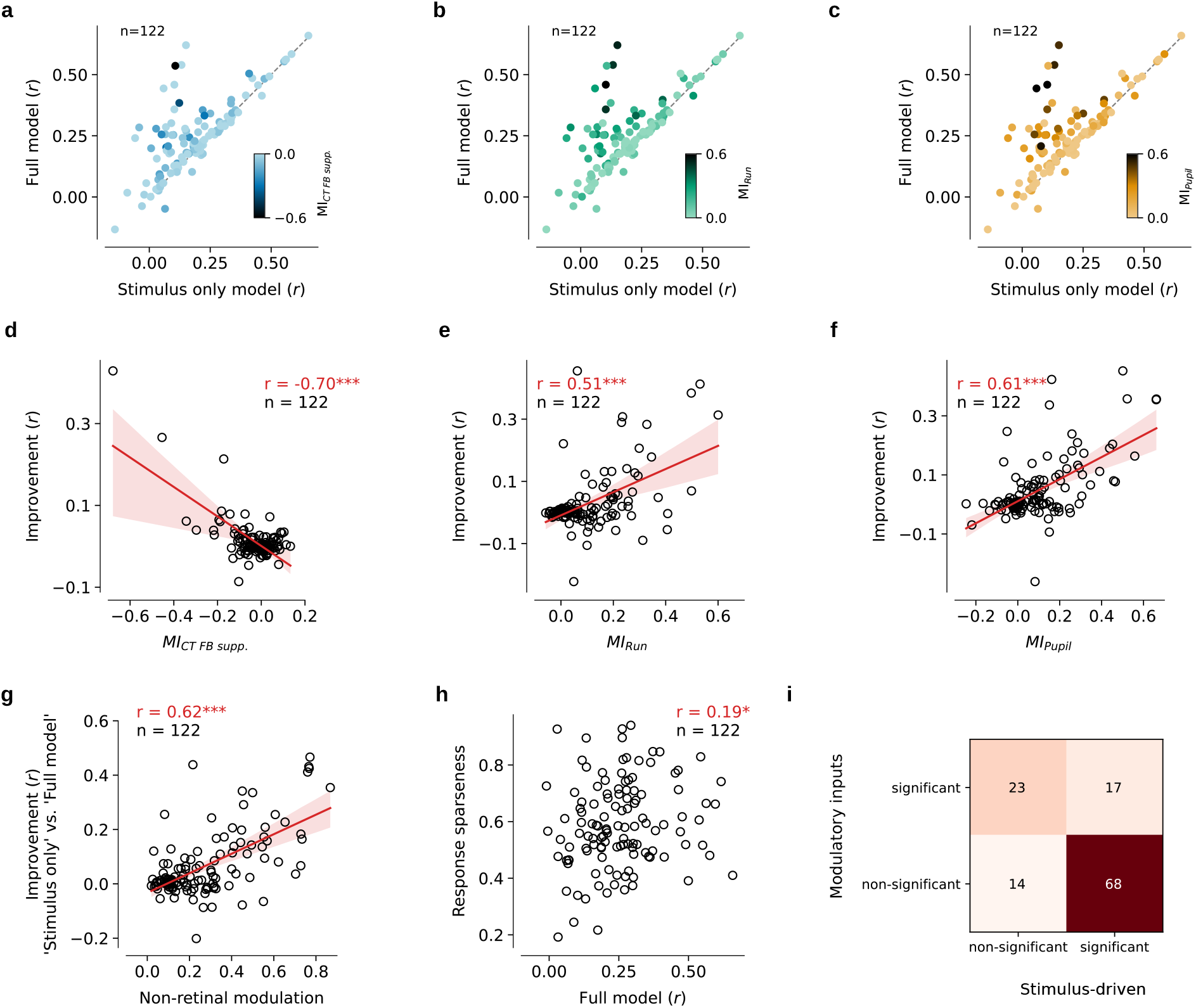
Model performance depends on various factors. (**a**) Comparison of model performances on the test set for all neurons in the ‘Stimulus only’ model and the ‘Full model’. Darker colours indicate stronger modulation by CT feedback quantified by *MI*_*CT FB supp*._ (see Methods). (**b**) Same as in (a) but for by running quantified by the modulation index *MI*_*Run*_. (**c**) Same as in (a) but for pupil size quantified by the modulation index *MI*_*Pupil*_. (**d**) Correlation of modulation by CT feedback suppression (*MI*_*CT FB supp*._) and model performance improvement. To measure improvement two models where compared: the ‘Stimulus only’ model and a model with Stimulus and CT feedback suppression as inputs. (**e**) Same as in h but for modulation by running (*MI*_*Run*_). The ‘Stimulus only’ model was compared with a model that integrated stimulus and running as inputs. (**f**) Same as in h but for modulation by pupil size (*MI*_*Pupil*_). The ‘Stimulus only’ model was compared with a model that integrated stimulus and pupil size as inputs. (**g**) Modulation by additional factors (*MI*_*Joint*_) versus improvement of model performance when comparing the ‘Stimulus only’ model and the ‘Full model’. * * \**p* ≤ 0.001 (**h**) Model performance versus response sparseness. Sparseness was computed according to (*112*). \**p* ≤ 0.05 (**i**) The matrix illustrates the neuron counts categorised into 68 neurons significant only to feedforward stimulus-related predictor, 23 neurons significant only to one or more of the predictors for modulatory inputs (CT feedback, run speed, and pupil dilation), 17 significant to all predictors, and 14 not exhibiting significance for any of the predictors.

**Figure S8.**
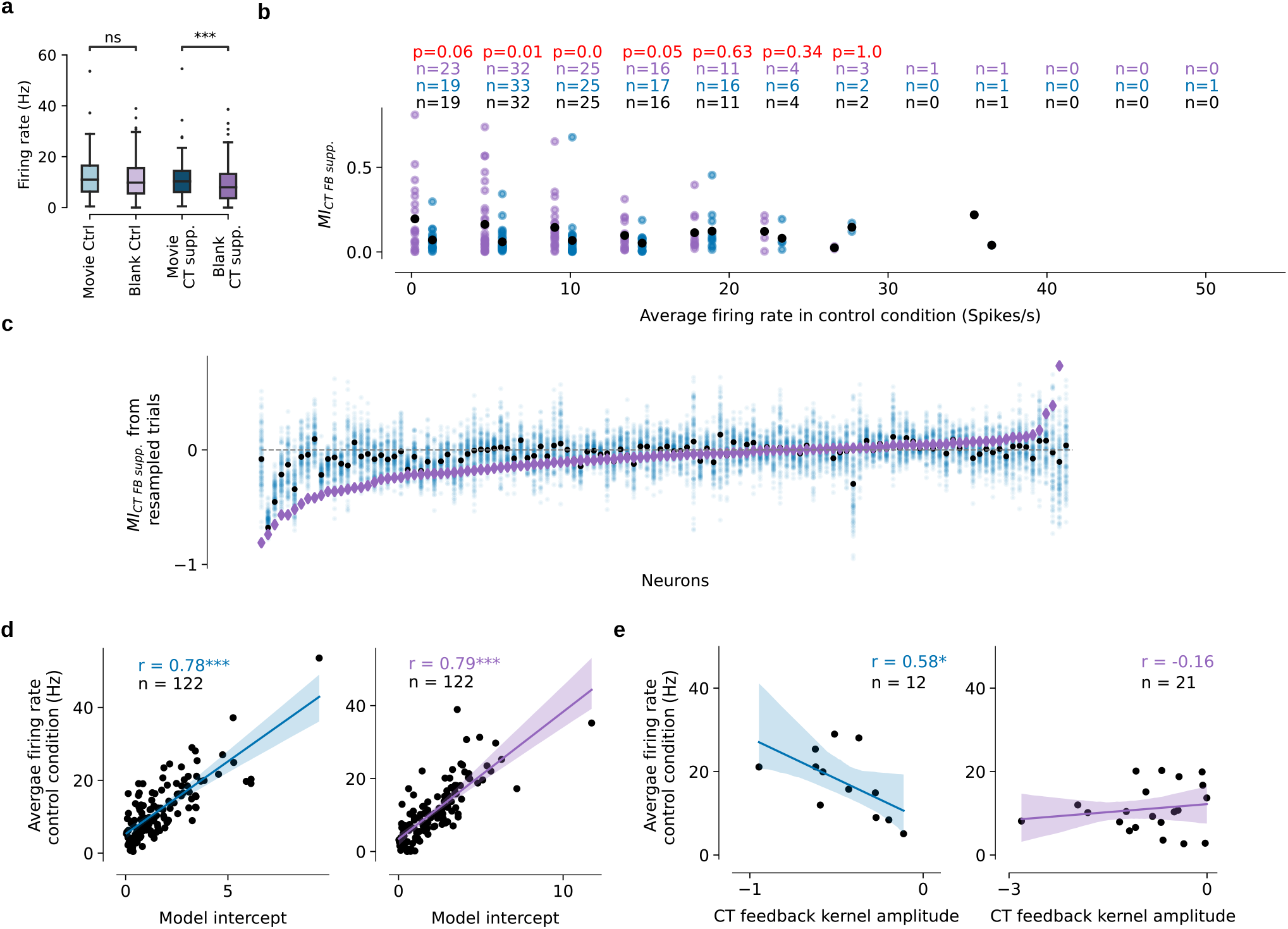
Influences of visual stimulus on effects of CT FB suppression in dLGN. (**a**) Firing rate comparison between conditions. Bonferroni corrected *p*-values of paired Wilcoxon signed-rank test: * * \**p* ≤ 1.0 × 10^−3^; ‘ns’ non-significant. (**b**) Analysis controlling potential influences of overall firing rate differences between conditions. *MI*_*CT FB supp*._ for movies (*blue*) or blank periods (*purple*) as a function of binned average firing rate during the control condition. We counted the neurons that fell within each firing rate bin for the two stimulus conditions (*top, coloured numbers*). To ensure equal number of neurons per firing rate bin between the conditions, we randomly sampled the minimum number of neurons for that bin for one of the stimulus conditions (*black numbers*). *Black dots* represent the mean *MI*_*CT FB supp*._ value per bin per condition after potential re-sampling. Even after approximating overall firing rates, *MI*_*CT FB supp*._ remained significantly higher for low firing rate bins, where most neurons were sampled (*red numbers*, Mann-Whitney-U test). (**c**) Scatters of *MI*_*CT FB supp*._ values recalculated based on re-sampled trials. Given that movie and blank conditions had different number of trials, and in order to rule out that the number of trials might affect the *MI*_*CT FB supp*._ values and accordingly the reported results, we performed 100 repeats of a re-sampling procedure. During each repeat, we randomly sampled 24 trials for each neuron during movie condition and re-calculated the *MI*_*CT FB supp*._ values (*light blue dots*). We found that the original movie *MI*_*CT FB supp*._ values (*black dots*) well represented the median of the re-calculated values. We also found that our observation that the original Blank *MI*_*CT FB supp*._ (*purple*) being stronger than movie *MI*_*CT FB supp*._ held true after re-sampling. (**d**) Scatter plot of average firing rate during the control condition vs. intercept fitted by the model for the movie (*left*) and blank periods (*right*). Panels (a–d) show data from all 122 neurons. (**e**) Scatter plot of average firing rate vs. maximum amplitude of the fitted CT feedback kernels for significantly modulated neurons for the movie (*left*) and blank (*right*) conditions.

